# RNA exosome-mediated RNA surveillance governs developmental timing in the human cerebellum

**DOI:** 10.64898/2026.04.19.719460

**Authors:** Nina A. Barr, Bozhidar Baltov, Rylee E. Kang, Jash J. Gada, Matthew J. Wade, Ethan N. Tjoa, Vivian Lee, Anoothi Seth, Kafui Dzirasa, Ashleigh E. Schaffer, Hyunmin Kim, Derrick J. Morton

## Abstract

Defects in RNA metabolism contribute to neurodevelopmental disease, yet how RNA surveillance shapes human brain development remains poorly understood. Mutations in genes encoding ubiquitously expressed RNA exosome subunits cause selective neurological disease, highlighting a poorly understood role for RNA surveillance in nervous system development. Here, we demonstrate that RNA exosome dysfunction alters the temporal coordination of gene expression during human cerebellar differentiation. Using CRISPR-engineered human cerebellar organoids modeling pathogenic EXOSC3 variants, we show that cerebellar lineages remain specified, but exhibit altered developmental trajectories, cellular composition, and maturation-associated transcriptional programs. Lineage-resolved transcriptomic and regulon analyses reveal selective disruption of developmental regulatory programs, particularly in Purkinje cells. These molecular abnormalities are accompanied by disorganized cerebellar architecture and impaired coordination of spontaneous neuronal activity. Together, our findings identify RNA exosome surveillance as a contributor to the temporal regulation of cerebellar development and support an early neurodevelopmental component of pontocerebellar hypoplasia type 1b pathogenesis.

## Introduction

Accurate RNA surveillance is fundamental to achieving transcriptome homeostasis during cellular differentiation^1, 2^. As cells transition between developmental states, extensive transcriptome remodeling generates nascent, unstable, and non-coding infrastructural and regulatory RNAs whose abundance must be precisely controlled to establish and resolve developmental gene expression programs^3^. While transcriptional control of cell fate decisions has been extensively characterized^4^, how RNA synthesis and decay are coordinated to regulate developmental transitions remains poorly understood^5, 6^.

A central regulator of post-transcriptional RNA surveillance is the RNA exosome, a conserved multiprotein ribonuclease complex that mediates 3’-5’ RNA processing and degradation of diverse classes of RNA, including coding, non-coding, and aberrant transcripts^7–11^. The RNA exosome comprises a catalytically inert nine-subunit core (EXOSC1-9) associated with ribonucleases DIS3 and EXOSC10^12^ (**Fig. 1A**); ribonuclease activities are further refined by adaptor and cofactor proteins that confer substrate specificity^13, 14^. Beyond the canonical roles of the complex in RNA quality control and rRNA processing^9, 15^, emerging evidence suggests that the RNA exosome regulates gene expression programs associated with cellular differentiation (reviewed in^16^).

**Figure 1.**
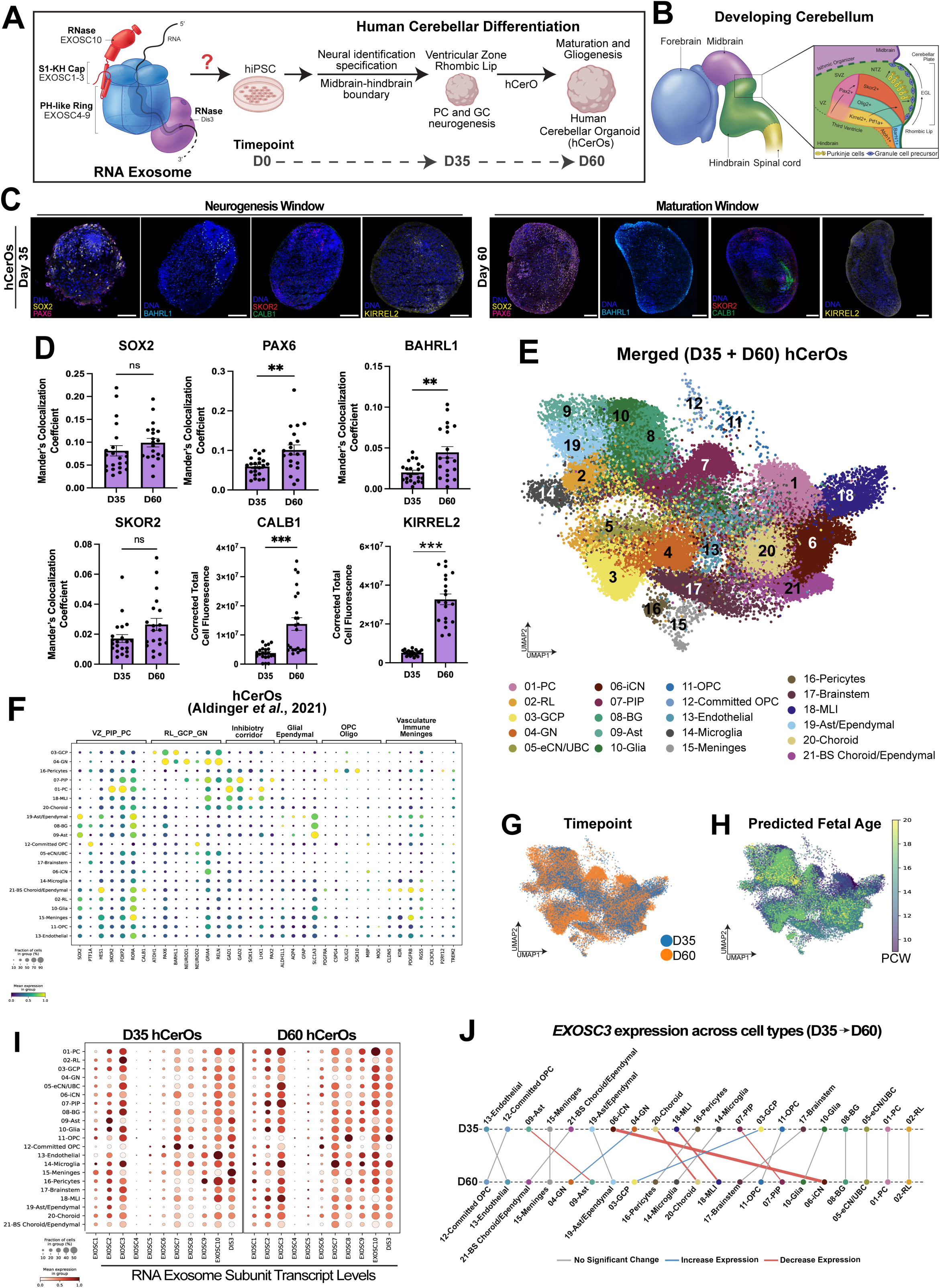
Human cerebellar organoids model cerebellar lineage specification and RNA exosome expression. **(A)** Structural representation of the RNA exosome and schematic of the human cerebellar organoid (hCerO) differentiation strategy from human induced pluripotent stem cells (hiPSCs). Organoids were analyzed at D35, corresponding to an active neurogenic stage, and D60, corresponding to early neuronal maturation. **(B)** Schematic of human cerebellar development illustrating ventricular zone (VZ)- and rhombic lip (RL)-derived lineages that generate major cerebellar neuronal and glial populations. **(C)** Representative immunofluorescence images of hCerOs at D35 and D60. D35 organoids contain SOX2+/KIRREL2 ventricular progenitors, PAX6+ rhombic lip domains, BARHL1+ granule precursors, SKOR2+ early Purkinje cells, and KIRREL2+rhombic lip domains. D60 organoids increased CALB1 expression consistent with Purkinje cell maturation. Nuclei are labeled with Hoechst. Scale bars, 200 μm. **(D)** Quantification of SOX2, PAX6, BARHL1, SKOR2, CALB1, and KIRREL2 immunofluorescence at D35 and D60. Each point represents the average expression across cryosectioned organoids. Colocalization of nuclei (Hoechst) and nuclear stains (SOX2, PAX6, BARHL1, SKOR2). Corrected total cell fluorescence was utilized for cytoplasmic stains (CALB1 and KIRREL2). Bars represent mean ± SEM. Statistical significance was determined using a Welch’s t-test; ns, not significant; **p<0.001; ***p<0.001. **(E)** UMAP projection of integrated single-cell RNA-seq profiles from D35 and D60 hCerOs identifying 21 cerebellar and associated cell populations. Cell identities are indicated by color. **(F)** Dot plot comparing canonical cell type marker expression in hCerOs with corresponding populations from the developing human fetal cerebellum atlas^69^. Dot size represents the fraction of cells expressing each gene, and color represents mean expression. **(G)** UMAP projection colored by developmental time point, showing the distribution of D35 (blue) and D60 (orange) cells with the integrated hCerO landscape^69, 70^. **(H)** UMAP projection colored by predicted fetal age in post-conception weeks (PCW) following mapping to the developing human fetal cerebellar reference atlas ^69^. **(I)** Dot plots showing transcript levels of RNA exosome subunits among annotated hCerO cell populations at D35 and D60. Dot size represents fraction of cells expressing each transcript, and color represents mean expression. **(J)** Line plot showing changes in EXOSC3 expression from D35 to D60 among annotated cell populations. For each cell type, differential *EXOSC3* expression between D35 and D60 was evaluated using the Wilcoxon rank-sum test in WT hCerOs. Changes with p<0.05 were considered statistically significant. Significant increases in expression are indicated by blue lines, significant decreases by red lines, and non-significant changes by gray lines. Line widths reflect the strength of the statistical significance, with thicker lines corresponding to smaller p values.

In pluripotent^17, 18^, epidermal^19, 20^, and hematopoietic^21–24^ contexts, RNA exosome-mediated decay regulates the abundance of lineage-specific transcripts and contributes to transitions between cellular identities^21^. Consistent with a role in cellular identity regulation, elevated expression of RNA exosome components has been associated with pluripotency, whereas reduced expression of certain components accompanies differentiation in some cellular systems^17, 25^. Together, these studies suggest that RNA exosome activity helps establish appropriate temporal boundaries between developmental gene expression programs, a role that remains poorly defined during neuronal development. More broadly, these findings raise the possibility that post-transcriptional RNA surveillance contributes to timing and coordination of developmental state transitions^26^. The RNA exosome is highly conserved in eukaryotes, ubiquitously expressed, and essential for viability in all systems examined^8, 27–30^. However, disruption of this broadly acting RNA surveillance complex produces striking tissue- and lineage-specific phenotypes, highlighting a central paradox in RNA biology^31^.

This paradox is particularly salient in the nervous system, where precise spatiotemporal control of RNA processing^32^, localization^33^, and stability^34^ is required for both development and long-term function. Neurons undergo extensive transcriptome remodeling during differentiation and maturation^35^, and defects in RNA metabolism are increasingly linked to neurodevelopmental and neurodegenerative disorders^36^. Mutations in genes encoding splicing factors, RNA binding proteins, helicases, and ribonucleases cause a spectrum of neurological diseases^36–40^, underscoring the heightened sensitivity of neural lineages to perturbations in intracellular RNA regulation. Why particular neuronal populations are disproportionately affected by dysfunction of broadly expressed RNA processing machinery remains poorly understood.

Autosomal recessive mutations in genes encoding structural subunits of the RNA exosome cause a spectrum of neurological disorders, including certain subtypes of Pontocerebellar Hypoplasia (PCH), a group of disorders characterized by impaired development and/or degeneration of the cerebellum, pons, and other neuronal populations^41^. Pontocerebellar Hypoplasia type 1b (PCH1b) is caused by biallelic variants in *EXOSC3*^42^, which encodes a structural subunit of the RNA exosome. PCH1b presents with profound cerebellar hypoplasia and progressive neurological dysfunction despite the ubiquitous requirement for RNA exosome activity^43–45^. How disruption of globally required RNA surveillance produces selective cerebellar vulnerability remains unresolved and represents a central question at the interface of RNA biology and neurodevelopment. Moreover, whether the cerebellar pathology associated with PCH1b reflects solely degeneration or originates, at least in part, from disruption of earlier developmental programs remains unclear.

The cerebellum offers a powerful system in which to define how RNA surveillance contributes to developmental state transitions. Cerebellar development requires the coordinated specification of ventricular zone (VZ)-derived inhibitory neurons and glia, and rhombic lip (RL)-derived excitatory neurons, followed by precisely ordered migration, lamination, and maturation^46–50^. Spatially patterned signaling cues and regionally restricted expression of developmental regulators establish distinct cerebellar progenitor domains and guide neuronal lineage specification^51, 52^. These processes depend on developmental gene expression programs that must be activated, remodeled, and resolved with lineage- and stage-specific precision^53, 54^. We therefore hypothesized that the RNA exosome contributes to the temporal coordination of cerebellar development by regulating lineage- and stage-specific gene expression programs. In this model, disruption of RNA surveillance would not necessarily prevent lineage specification but could alter the timing, magnitude, or cellular specificity of developmental programs as cells progress toward mature states. Such effects could provide a mechanism by which dysfunction of a ubiquitously expressed RNA processing complex produces selective vulnerability among cerebellar lineages.

Notably, mouse models of related RNA processing disorders, including CLP1-associated PCH^55, 56^, fail to reproduce key human cerebellar phenotypes despite clear molecular defects, highlighting a divergence in cerebellar vulnerability and developmental programs between species. To address this, we generated human cerebellar organoids (hCerOs) derived from human induced pluripotent stem cells (hiPSCs) as an experimentally tractable model of human cerebellar differentiation^57, 58^. Using this system, we interrogated two PCH1b disease-associated *EXOSC3* missense variants, Gly31Ala (G31A)^42, 59–63^ and Gly191Cys (G191C)^64^, in an isogenic genetic background. Integrating single-cell transcriptomics, developmental trajectory analysis, lineage-resolved transcriptional and regulon analysis, spatial characterization, and functional recordings of neuronal activity, we determined how RNA exosome dysfunction alters human cerebellar development over time.

We find that major cerebellar lineages remain specified and continue to progress through developmental time following EXOSC3 disruption, while their developmental trajectories, relative abundance, spatial organization, and maturation-associated transcriptional programs are altered. These effects are graded between EXOSC3 variants and disproportionately affect specific neuronal populations, particularly the Purkinje cell lineage. Regulon analysis further reveals selective disruption of lineage-associated regulatory programs despite ubiquitous RNA exosome expression, providing a framework for cell type-specific developmental vulnerability. These abnormalities are accompanied by disrupted cerebellar tissue organization and reduced coordination of spontaneous neuronal activity. Together, our findings identify RNA exosome-mediated surveillance as an important contributor to the temporal coordination of human cerebellar development and support an early neurodevelopmental component of PCH1b pathogenesis.

## Results

### Human cerebellar organoids recapitulate lineage diversity and broad RNA exosome expression

To investigate how RNA exosome dysfunction influences human cerebellar development, we generated a 3D human cerebellar organoid (hCerO) platform that recapitulates key stages of cerebellar lineage specification (**Fig. 1A**). Isogenic hiPSCs were differentiated through neural induction and midbrain-hindbrain boundary specification into ventricular zone (VZ) and rhombic lip (RL) progenitor domains that give rise to Purkinje cells and granule neurons, respectively (**Fig. 1B**). Organoids were analyzed at day 35 (D35), representing an early neurogenesis window, and day 60 (D60), representing a maturation window.

Immunofluorescence analyses confirm the establishment of major cerebellar progenitor and neuronal populations during each developmental stage (**Fig. 1C**). At D35, hCerOs contain SOX2+ neural progenitors, PAX6+ rhombic lip (RL) derivatives, and BARHL1+ granule cell precursors. The ventricular zone (VZ) derivatives: KIRREL2+ cell adhesion marker, SKOR2+ early Purkinje cells, and comparatively low levels of the Purkinje maturation marker CALB1. SOX2+ and PAX6+ populations occupy partially distinct but intermingled regions, consistent with spatial compartmentalization characteristic of neural organoids^65, 66^. By D60, these lineage-associated populations remain detectable while CALB1 expression increased, consistent with continued progression of the Purkinje lineage toward a more mature state (**Fig. 1C**). Quantification of lineage-marker levels confirms developmental changes between D35 and D60 (**Fig. 1D**).

To define cellular diversity of hCerOs at single-cell resolution, we performed single-cell RNA sequencing (scRNA-seq) on pooled D35 and D60 organoids from two biological replicates per time point, with 8-12 organoids pooled per clonal replicate and approximately 8,000 cells analyzed per sample. hCerOs were dissociated, fixed, permeabilized, and barcoded prior to sequencing on the Novaseq X platform^67, 68^. Quality-control analysis demonstrates comparable relationships between transcript counts and detected genes in all biological replicates at both developmental stages, with low mitochondrial transcript content (**Fig. S1**). Integrated analysis identified major cerebellar populations, including Purkinje cells (PC), granule cell precursors (GCP), granule neurons (GN), interneurons (iCN), astroglia (Ast), oligodendrocyte precursor cells (OPC), Bergmann glia (BG), and additional neural and non-neural populations (**Fig. 1E**). Comparison with the Aldinger fetal cerebellar atlas^69^ confirms expression of lineage-associated transcriptional programs corresponding to VZ-derived inhibitory, RL-derived excitatory, glial, and additional cerebellar populations **(Fig. 1F**). Having established correspondence with the developing human cerebellum, we next examined developmental stage within the hCerO dataset. D35 and D60 cells show distinct but overlapping distributions within the integrated embedding (**Fig. 1G**), while projection of predicted fetal age positions hCerO populations along developmental states corresponding to approximately 14-24 post-conception weeks (PCW) (**Fig. 1H**).

To further assess the correspondence of hCerOs with independent developmental references, we integrated our dataset with the Aldinger fetal cerebellar atlas^69^ and the Human Neural Organoid Cell Atlas (HNOCA)^70^, using an scVI-based framework (**Fig. S1A-B**). Joint embedding reveals dataset-dependent separation rather than complete transcriptional overlap, with hCerOs showing greater correspondence to the fetal cerebellar reference than to the broader HNOCA reference (**Fig. S1A-B**). Prediction entropy varied across clusters and reference datasets, with several hCerO populations showing lower-entropy assignments to the Aldinger reference and less confident correspondence to HNCOA (**Fig. S1C-D**). Detailed cross-reference mapping further shows that hCerOs contain early cerebellar populations that are incompletely represented in the broader neural organoid reference (**Fig. S1E**). Thus, external benchmarking supports the cerebellar identity of the major hCerO populations while also highlighting transcriptional differences between this model and existing fetal and organoid reference datasets.

We next examined expression of RNA exosome components of the identified cerebellar populations. Core RNA exosome subunit transcripts are broadly detected across progenitor, neuronal, and glial populations at both D35 and D60 (**Fig. 1I**). We then examined developmental changes in *EXOSC3* expression within individual cell types (**Fig. 1J**). *EXOSC3* remains broadly expressed at both developmental stages, but the magnitude and direction of expression changes between D35 and D60 varied by cell type. Several populations show little change in *EXOSC3* expression, whereas others exhibit increased or decreased expression during maturation (**Fig. 1J**). Thus, *EXOSC3* is not restricted to a particular cerebellar lineage, but its expression is differentially regulated during development in a cell type-dependent manner.

Together, these findings establish that hCerOs recapitulates key features of early human cerebellar development, including the generation of major VZ- and RL-derived populations, cellular diversity, and progression across developmental states. Comparison with fetal cerebellar and organoid reference datasets further supports the identity of the major cerebellar populations while defining features unique to the hCerO model. Broad expression of RNA exosome components in these developmental populations provides a foundation for determining how pathogenic EXOSC3 variants alter cerebellar lineage progression.

### *EXOSC3* mutations alter cerebellar lineage progression in human cerebellar organoids

To determine how RNA exosome dysfunction affects human cerebellar lineage progression, we modeled two pathogenic EXOSC3 variants associated with Pontocerebellar Hypoplasia Type 1b: G31A^42^ and G191C^64^, alongside an isogenic wildtype control (EXOSC3^WT^) (**Fig. 2A**). Using CRISPR/Cas9 editing, we introduced each variant into hiPSCs and differentiated the resulting lines into human cerebellar organoids (hCerOs) utilizing an established protocol^57, 58^ **(Fig. 2A**). Previous studies in *Drosophila* established that the EXOSC3^G31A^ equivalent variant (Rrp40^G11A^) produces more pronounced defects in RNA exosome function than the EXOSC3^G191C^ equivalent (Rrp40^G146C^), as evidenced by differences in steady-state transcript abundance and rRNA processing^27, 28^. These findings support a spectrum of RNA exosome dysfunction and provide rationale for investigating shared and variant-specific effects on human cerebellar lineage progression and maturation.

**Figure 2.**
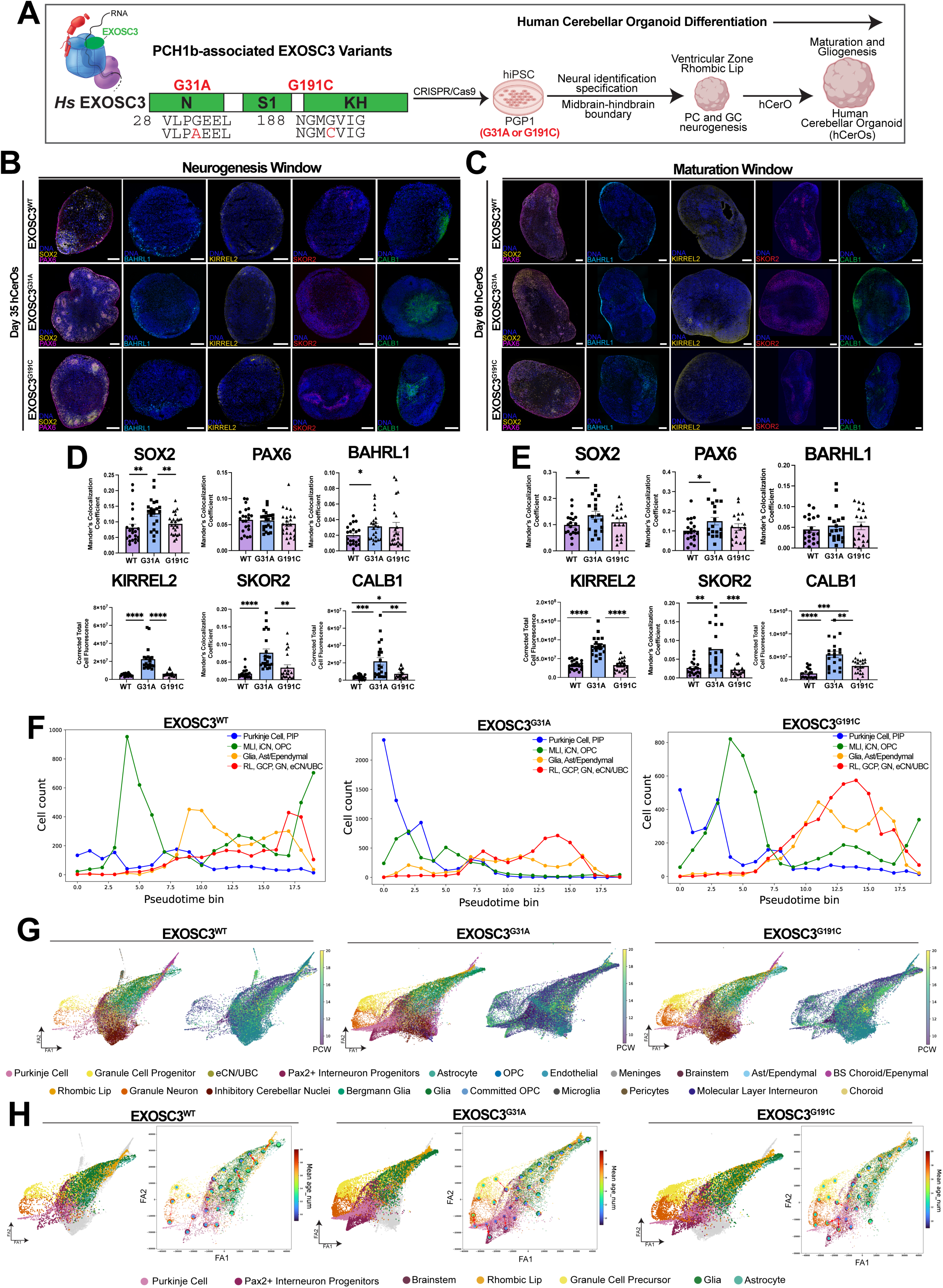
*EXOSC3* mutations impair temporal resolution of cerebellar lineage progression. **(A)** Schematic of the PCH1b-associated G31A and G191C variants within the human RNA exosome cap subunit EXOSC3 and experimental strategy for generating CRISPR-engineered isogenic hiPSCs and differentiating them into human cerebellar organoids (hCerOs). Organoids were analyzed at D35 during neurogenesis and D60 during early maturation **(B)** Representative immunofluorescence images of EXOSC3^WT^, EXOSC3^G31A^, and EXOSC3^G191C^ hCerOs at D35. SOX2 and KIRREL2 label ventricular zone progenitors; PAX6 and KIRREL2 identify rhombic lip-associated domains; BAHRL1 labels granule lineage cells; SKOR2 labels early Purkinje lineage cells; and CALB1 marks Purkinje cell maturation. Nuclei were labeled with Hoechst. Scale bar, 200 μm **(C)** Representative immunofluorescence images of EXOSC3^WT^, EXOSC3^G31A^, and EXOSC3^G191C^ hCerOs at D60 showing genotype-dependent differences in cerebellar lineage organization and maturation. Markers are as described in (B). Nuclei were labeled with Hoechst. Scale bar, 200 μm **(D)** Quantification of SOX2, PAX6, BARHL1, KIRREL2, SKOR2, and CALB1 immunofluorescence in D35 hCerOs. Each point represents an independent organoid. Bars represent mean ± SEM. Statistical significance was determined using Welch’s t-test; ns, not significant;, *p<0.05;, **p<0.01;, ***p<0.001; ****p<0.0001. **(E)** Quantification of SOX2, PAX6, BARHL1, KIRREL2, SKOR2, and CALB1 immunofluorescence in D60 hCerOs. Data are presented and analyzed as described in (D). **(F)** Distribution of major cerebellar lineages along pseudotime for EXOSC3^WT^, EXOSC3^G31A^, and EXOSC3^G191C^ hCerOs. Fine-grained cell identities were grouped into Purkinje lineage (PC/PIP), inhibitory lineage (MLO/iCN/OPC), glial lineage (Glia/Ast/Ependymal), and rhombic lip-derived lineage (RL/GCP/GN/eCN/UBC). Cell counts within pseudotime bins illustrate genotype-dependent differences in the distribution of cerebellar lineages along the inferred developmental trajectory. **(G)** Developmental trajectory embeddings for EXOSC3^WT^, EXOSC3^G31A^, and EXOSC3^G191C^ hCerOs. For each genotype, the left embedding is colored by annotated cell identity and the right by predicted developmental age in post-conception weeks (PCW), illustrating the relationship between lineage identity and developmental progression. **(H)** Centroid-based ForceAtlas (FA) trajectory maps summarizing developmental relationships between lineages. Nodes represent lineage centroids (outer ring: cell type composition; inner circle: mean developmental age), connected by minimum spanning trees with arrows indicating increasing post-conception weeks (PCW). EXOSC3^WT^ organoids display coherent trajectories linking ventricular zone-derived inhibitory lineages (Purkinje cells (PC) and Pax2+ interneuron progenitors (PIP)) and rhombic lip-derived excitatory transitions (RL -> GCP -> GN), whereas *EXOSC3* mutants exhibit disrupted network organization and mixed developmental states.

Sanger sequencing confirms precise editing at the *EXOSC3* locus in hiPSCs (**Fig. S2A-B**). To assess the reproducibility of genotype-dependent developmental programs, we independently analyzed scRNA-seq datasets generated from two distinct hiPSC clones per genotype. Replicate-resolved expression trajectories reveal broadly concordant developmental patterns for cerebellar lineage markers, including SOX2, KIRREL2, PAX6, BAHRL1, and SKOR2, supporting the reproducibility of genotype-associated transcriptomic changes across independent clones (**Fig. S2C**). Organoids from all genotypes form successfully, although genotype-dependent differences in growth are observed across differentiation (**Fig. S3A**), indicating that EXOSC3 variants alter organoid growth without preventing cerebellar organoid formation. Mitochondrial transcript content remained below 5% in all genotypes **(Fig. S3I**), indicating comparable mitochondrial RNA fractions and no genotype-dependent differences in this single-cell RNA-seq quality control metric.

We first examined the consequences of EXOSC3 dysfunction during the neurogenesis window (D35). EXOSC3^WT^ hCerOs establish cerebellar progenitor and neuronal populations labeled by neuronal progenitor, SOX2, the rhombic lip-associated markers PAX6 and BARHL1, ventricular zone cell adhesion marker KIRREL2, and the early Purkinje cell marker SKOR2, with comparatively low expression of the Purkinje maturation marker CALB1 (**Fig. 2B, D**). *EXOSC3* mutant organoids also generate these populations, indicating that the variants do not prevent initial cerebellar lineage specification. However, EXOSC3^G31A^ organoids exhibit increased SOX2, BARH1, and KIRREL2 labeling at D35, consistent with increased representation of progenitor and rhombic lip-associated programs during the neurogenic window (**Fig. 2D**). Changes in PAX6, SKOR2, and CALB1 are comparatively modest and varied between the two mutant genotypes. Thus, EXOSC3 dysfunction permits establishment of major cerebellar lineages but alters the relative representation of early developmental programs during neurogenesis.

We next examined lineage-associated markers during the maturation window (D60) (**Fig. 2C**, **2E**). EXOSC3^WT^ hCerOs exhibit increased CALB1 expression, consistent with progression of the Purkinje lineage toward a more mature state, together with developmental changes in progenitor and rhombic lip-associated markers. *EXOSC3* mutant organoids similarly retain the major cerebellar populations and express CALB1, indicating that maturation-associated programs are initiated rather than completely blocked. However, EXOSC3^G31A^ organoids exhibit increased PAX6, KIRREL2, and SKOR2 labeling at D60, indicating increased representation of progenitor, rhombic lip-associated, and early Purkinje lineage programs during the maturation window (**Fig. 2E**). CALB1 is also altered in the mutant organoids, demonstrating that these early lineage-associated differences occur alongside, rather than instead of, activation of the Purkinje maturation program. EXOSC3^G191C^ organoids show a more restricted pattern of marker changes at D60. Thus, EXOSC3 dysfunction does not prevent cerebellar lineage specification or the initiation of maturation but alters the relative representation and coordination of early and maturation-associated lineage programs.

Given that the altered representation of lineage-associated markers could reflect differences in progenitor proliferation, we next examined cell proliferation across developmental stages using pulse-chase BrdU labeling. Proliferation is significantly increased in *EXOSC3* mutant organoids at D35, whereas no significant difference is observed at D60 (**Fig. S3C-D**). Thus, EXOSC3 dysfunction produces a stage-specific increase in proliferation during early neurogenesis that is not maintained during the maturation window. This transient proliferative phenotype may contribute to the altered representation of early lineage-associated population observed at D35 but does not account for the lineage differences that remain evident at later developmental stages. We therefore leveraged single-cell transcriptomic analyses to determine whether EXOSC3 dysfunction alters the temporal progression of cerebellar lineage beyond proliferation,

To further assess developmental progression, we examined the distribution of cerebellar lineages across pseudotime bins (**Fig. 2F**). In EXOSC3^WT^ organoids, cerebellar populations show distinct distributions across pseudotime, with the Purkinje/PAX2+ interneuron progenitor population concentrated toward earlier developmental intervals and rhombic lip-derived neuronal and glial populations distributed across later intervals. In contrast, *EXOSC3* mutant organoids show redistribution of these populations across pseudotime, with the most pronounced differences observed in EXOSC3^G31A^ organoids. In particular, Purkinje/PAX2+ interneuron progenitor populations are enriched at earlier pseudotime positions in EXOSC3^G31A^ organoids, while rhombic lip-derived and glial populations exhibit altered distributions across later pseudotime intervals. EXOSC3^G191C^ organoids show a more intermediate pattern, retaining greater similarity to the wild-type distribution while exhibiting shifts in several lineage populations. Together, these findings indicate that RNA exosome dysfunction alters the temporal distribution of cerebellar cell states rather than producing a uniform delay in differentiation.

To quantify maturation in control and mutant hCerOs, we leveraged our single-cell RNA-seq data to examine lineage-aggregated cell counts across pseudotime bins (**Fig. 2G**). EXOSC3^WT^ organoids show the expected developmental progression, characterized by a decline in progenitor states and the resolution of rhombic lip-derived populations, concomitant with the emergence of mature Purkinje cells. In contrast, mutant organoids accumulate early Purkinje cell states that remain intermingled with mature Purkinje populations, indicating failure to resolve developmental identities. This phenotype is most pronounced in EXOSC3^G31A^, where early-born Purkinje cells persist across pseudotime, whereas EXOSC3^G191C^ shows partial resolution toward a wildtype distribution. Among all cerebellar lineages, Purkinje cells exhibit the most pronounced developmental disruption, consistent with their known vulnerability in PCH1b^64^.

To further resolve how EXOSC3 dysfunction affects developmental progression, we projected cerebellar cell identities and inferred developmental age onto the developmental landscape (**Fig. 2G**). EXOSC3^WT^ organoids exhibited structured relationships among cerebellar populations and developmental age, with Purkinje/PAX2+ interneuron progenitors and rhombic lip-derived populations occupying distinct regions of the developmental landscape. In *EXOSC3* mutant organoids, the distribution of cell identities relative to developmental age was altered, indicating that lineage identity and developmental progression become less tightly coordinated following RNA exosome dysfunction. These differences were most apparent in EXOSC3^G31A^ organoids, whereas EXOSC3^G191C^ retained greater similarity to the wild-type developmental organization. Together with the pseudotime redistribution observed in **Fig. 2F**, these findings indicate that EXOSC3 dysfunction alters the temporal organization of cerebellar lineage progression rather than uniformly arresting differentiation.

To capture these developmental disruptions, we summarized the data using a centroid-based representation in the same FA embedding space, in which centroids encode cell type composition (outer ring) and mean developmental age (inner ring) and are connected by a minimum spanning tree oriented from lower to higher post-conception weeks (PCW) (**Fig. 2H**). This representation provides an interpretable approximation of developmental state relationships. In EXOSC3^WT^ organoids, major cerebellar populations occupy distinct positions across the developmental landscape, including ventricular zone-derived inhibitory populations comprising Purkinje cells (PCs) and PAX2+ interneuron progenitors (PIPs), as well as rhombic lip-derived excitatory populations spanning rhombic lip (RL) granule cell precursor (GCP), and granule (GN) states. In contrast, *EXOSC3* mutant organoids exhibit altered relationships between cell identity and developmental age across this landscape, with particularly prominent changes among Purkinje-associated and rhombic lip-derived populations. These alterations are most pronounced in EXOSC3^G31A^ organoids, whereas EXOSC3^G191C^ retains greater similarity to the wildtype developmental organization. Together, these findings indicate that RNA exosome dysfunction alters the temporal organization and coordination of cerebellar lineage progression rather than producing a uniform developmental arrest.

Together, these findings demonstrate that pathogenic EXOCS3 variants permit the establishment of major cerebellar lineages but alter their progression across developmental states. Rather than producing a uniform developmental arrest, RNA exosome dysfunction changes the timing and distribution of lineage progression, with the strongest perturbations observed in EXOSC3^G31A^ organoids. We next asked whether these altered developmental trajectories are accompanied by changes in cerebellar tissue organization and lineage composition during maturation.

### EXOSC3 mutations disrupt cerebellar tissue organization and lineage composition during maturation

We next asked whether altered developmental progression in *EXOSC3* mutant hCerOs is accompanied by changes in cerebellar tissue organization and lineage composition. At D60, EXOSC3^WT^ organoids contain spatially organized BARHL1+ granule lineage cells and CALB1+ Purkinje cells, consistent with emergence of cerebellar-like laminar organization (**Fig. 3A**). EXOSC3^G31A^ and EXOSC3^G191C^ organoids generate both populations but exhibit altered spatial organization, including BARHL1+ cells positioned ectopically within or below CALB1+ Purkinje cell layer (**Fig. 3A**). Quantification shows no significant genotype-dependent differences in the overall position of the external granule or Purkinje cell layers, whereas ectopic BARHL1+ cells below the Purkinje layer are significantly increased in both EXOSC3 mutants (**Fig. 3B**). Thus, EXOSC3 dysfunction did not prevent formation or gross position of these cerebellar populations but disrupted their spatial organization during maturation (**Fig. 3C**).

**Figure 3.**
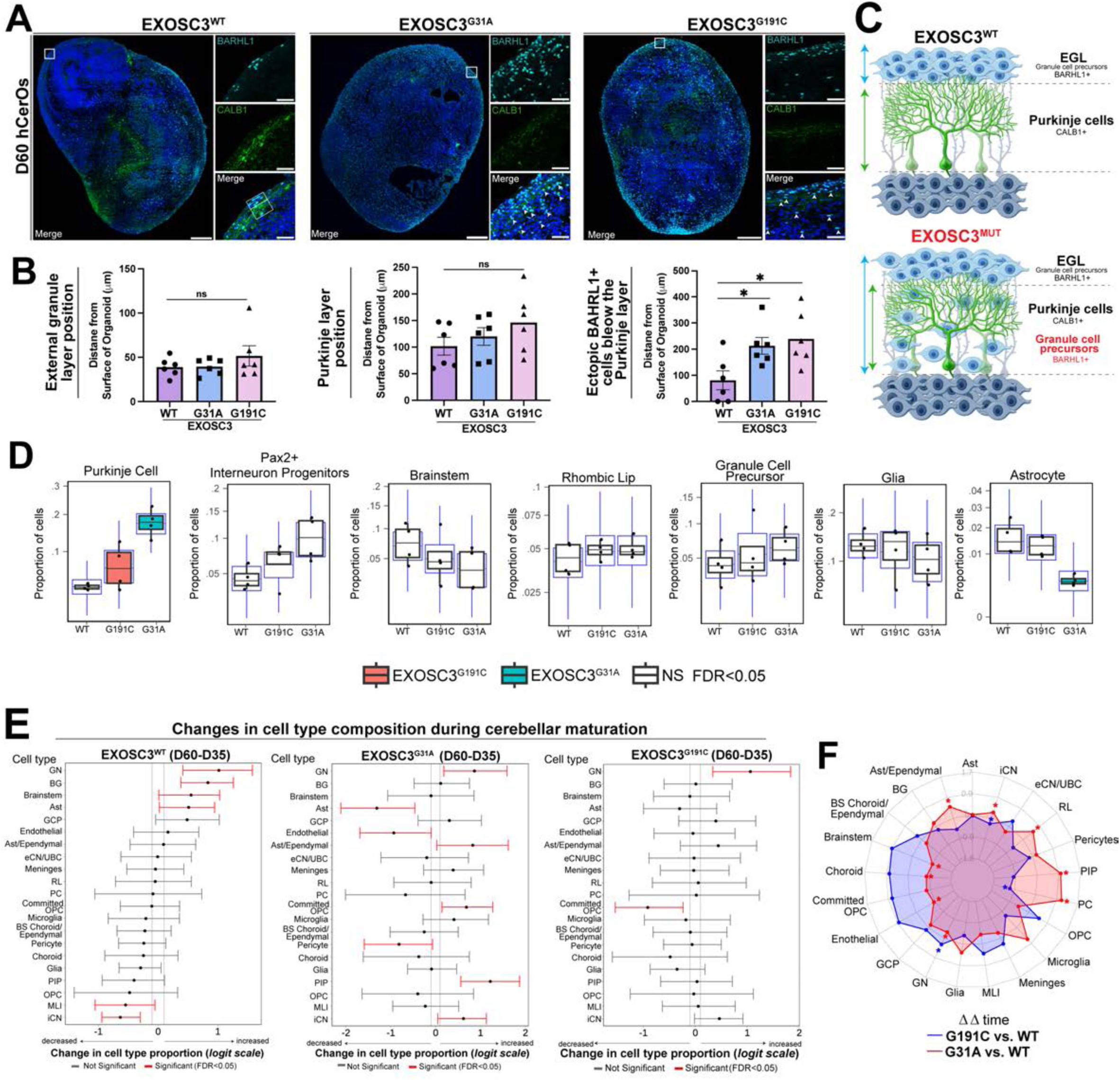
*EXOSC3* mutations disrupt cerebellar tissue organization and lineage composition during maturation. **(A)** Representative immunofluorescence images of D60 EXOSC3^WT^, EXOSC3^G31A^, EXOSC3^G191C^ human cerebellar organoids (hCerOs) stained for BARHL1 (granule cell lineage) and CALB1 (Purkinje cells). Insets show the spatial relationship between BARHL1+ and CALB1+ populations. Nuclei were labeled with Hoechst. Scale bars: Whole organoid: 200 µm (20X); Zoomed layering: 50 µm (20X). **(B)** Quantification of cerebellar tissue organization at D60. Left, position of the external granule layer (BARHL1+) relative to the organoid surface. Middle, position of the Purkinje cell layer (CALB1+) relative to the organoid surface. Right, number of ectopic BARHL1+ cells positioned within or below the CALB1+ Purkinje cell layer. Overall positioning of the external granule and Purkinje cell layers was not significantly different among genotypes, whereas ectopic BARHL1+ cells were increased in *EXOSC3* mutant hCerOs. Each point represents independent organoids; bars represent mean ± SEM. Statistical significance was determined using Welch’s t-test; ns, not significant; *p<0.05. **(C)** Schematic illustrating cerebellar-like organization in EXOSC3^WT^ and *EXOSC3* mutant hCerOs. In WT hCerOs, BARHL1+ granule-lineage cells within the external granule layer (EGL) and CALB1+ Purkinje cells occupy spatially organized domains. *EXOSC3* mutant hCerOs exhibit increased ectopic BARHL1+ cells within the region below the Purkinje cell layer. **(D)** Cell type composition in EXOSC3^WT^, EXOSC3^G31A^, and EXOSC3^G191C^ hCerOs for selected cerebellar populations, including Purkinje cells, PAX2+ interneuron progenitors, brainstem-associated cells, rhombic lip cells, granule cell precursors, glia, and astrocytes. Box plots show the proportion of cells assigned to each population. Significant genotype-dependent differences are indicated at FDR <0.05; ns, not significant. **(E)** Changes in cell type composition during D35-to-D60 cerebellar maturation for EXOSC3^WT^, EXOSC3^G31A^, EXOSC3^G191C^ hCerOs. Forest plots show estimated changes in cell type proportions on a logit scale with corresponding 95% confidence intervals. Significant developmental changes (FDR<0.05) are shown in red, and non-significant changes in black. **(F)** Relative genotype-dependent changes in cell type composition during maturation. Radar plot shows the difference in D35-to-D60 compositional change (ΔΔtime) for EXOSC3^G191C^ and EXOSC3^G31A^ relative to EXOSC3^WT^ among major cerebellar and associated cell populations. Values represent genotype-dependent deviations from the developmental change observed in WT, illustrating lineage-selective effects of EXOSC3 dysfunction during maturation.

We next determined whether these architectural abnormalities are accompanied by changes in the relative abundance of cerebellar cell populations. Single-cell analysis revealed lineage-selective differences in cell type composition rather than a uniform change in cellular abundance (**Fig. 3D**). Purkinje cells show the most prominent increase in EXOSC3^G31A^ organoids, with EXOSC3^G191C^ showing an intermediate phenotype. PAX2+ interneuron progenitors (PIPs) exhibit a similar genotype-associated increase. In contrast, brainstem-associated and astrocytic populations are reduced, particularly in EXOSC3^G31A^ organoids, whereas rhombic lip, granule cell precursor (GCP), and broader glial populations show comparatively modest differences (**Fig. 3D, Fig. S4**). Notably, the reduction in astrocytes argues against a generalized shift from neuronal toward glial identity as an explanation for the observed changes in neuronal populations.

Because the developmental trajectory analyses indicate that *EXOSC3* mutations alter progression through cerebellar cell states, we next examined how the relative abundance of individual populations changed between D35 and D60 within each genotype (**Fig. 3E**). EXOSC3^WT^ organoids show substantial developmental remodeling between D35 and D60, with increases in several neuronal and glial populations accompanied by decreases in earlier progenitor and intermediate populations. *EXOSC3* mutant organoids also show substantial compositional remodeling between D35 and D60, showing that differentiation continued despite RNA exosome dysfunction. However, the magnitude and distribution of these changes differed from EXOSC3^WT^ and between EXOSC3 variants (**Fig. 3E**). Comparing maturation-associated changes relative to EXOSC3^WT^ further reveals that EXOSC3^G31A^ and EXOSC3^G191C^ affected individual populations to different extents, with particularly prominent differences involving Purkinje cells and PAX2+ interneuron progenitors (PIPs) (**Fig. 3F**).

Together, these findings demonstrate that EXOSC3 dysfunction does not produce a uniform developmental arrest. Major cerebellar lineages are generated and continue to change in abundance during maturation, but their relative representation and spatial organization are altered in a genotype- and lineage-dependent manner. The accumulation of Purkinje-lineage populations, together with ectopic positioning of granule-lineage cells, supports a model in which impaired RNA surveillance disrupts the coordinated progression and organization of developing cerebellar lineages.

### EXOSC3 dysfunction disrupts the temporal coordination of neuronal network activity

To determine whether altered developmental progression in *EXOSC3* mutant hCerO is accompanied by functional deficits, we examined spontaneous neuronal activity in ∼D150 organoids using calcium imaging (**Fig. 4A)**. Spontaneous calcium transients are detected in EXOSC3^WT^, EXOSC3^G31A^, and EXOSC3^G191C^ hCerOs, indicating that neurons in each genotype acquired functional activity. However, EXOSC3^G31A^ organoids show a significant reduction in event frequency compared with EXOSC3^WT^, whereas EXOSC3^G191C^ shows a more modest reduction (**Fig. 4B**). In contrast, inter-spike intervals do not differ significantly among genotypes. Analysis of pooled single-cell calcium phenotypes further shows that active cells are present in all three genotypes, with variable effects on event amplitude, active-time fraction, event participation, and temporal regularity (**Fig. S5A**). Thus, EXOSC3 dysfunction alters the frequency of spontaneous neuronal activity without eliminating neuronal activity.

**Figure 4.**
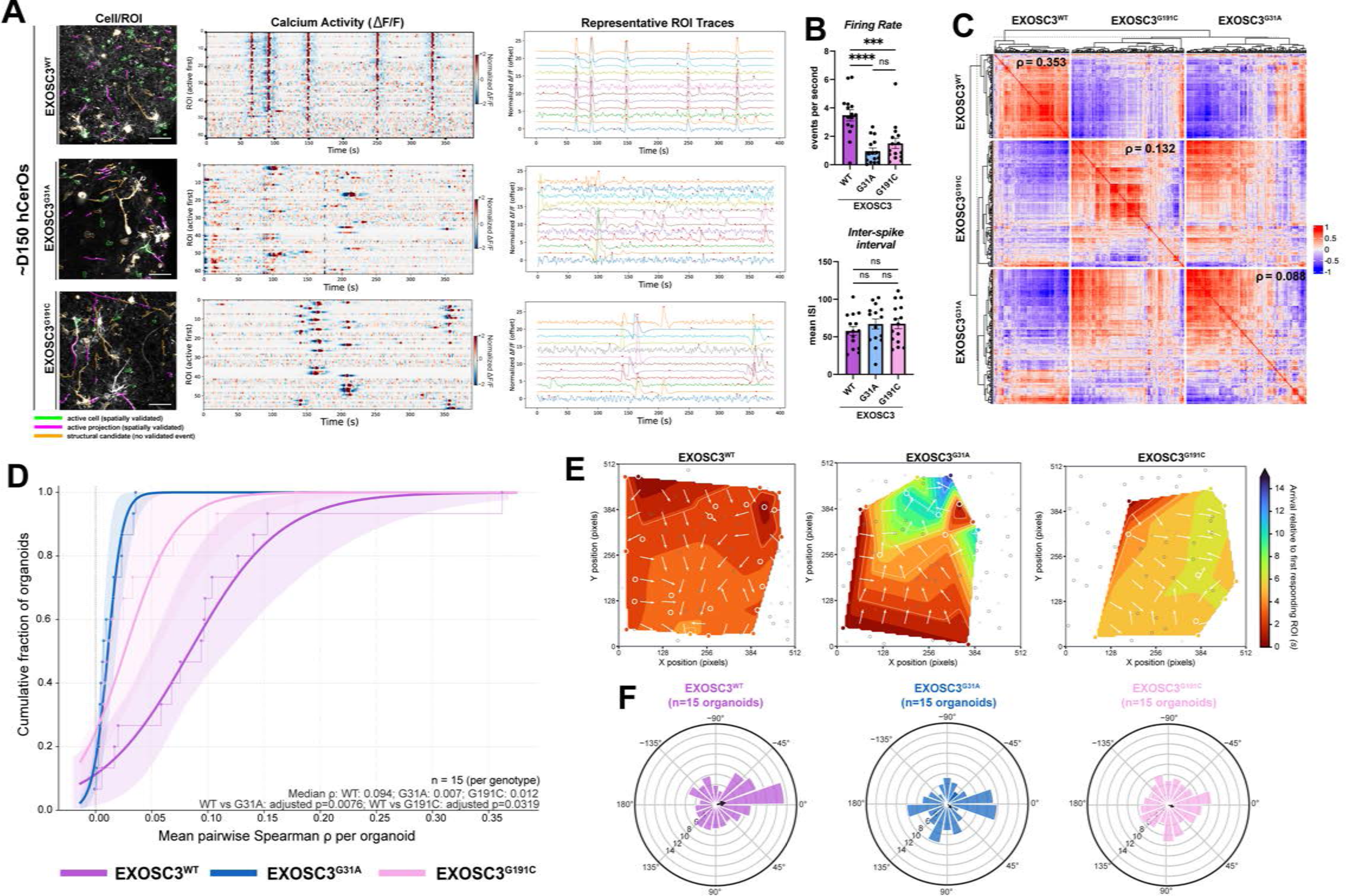
EXOSC3 variants disrupt the coordination of neuronal network activity. **(A)** Representative spontaneous calcium recordings from ∼D150 EXOSC3^WT^, EXOSC3^G31A^, EXOSC3^G191C^ human cerebellar organoids (hCerOs), arranged from top to bottom. For each genotype, the left panel shows identified regions of interest (ROIs) contours; the middle panel shows normalized ΔF/F over time, with one row per ROI and event-positive ROIs placed first; and the right panel shows selected ROI traces from the same recording, vertically offset for visibility. The horizontal axes show elapsed time in seconds across approximately 390s of imaging (1.18 frames/s). Early near-vertical bands in the WT heatmap indicate widespread, temporally coordinated calcium transients. The heatmap alone does not establish a propagating wave; spatial recruitment is assessed in panel E. Occasional aligned responses in G191C indicate transient coactivation of subsets of ROIs rather than WT-like population-wide recruitment. Scale bar, 200 μm. **(B)** Quantification of spontaneous calcium-event frequency (top) and mean inter-spike interval (bottom). EXOSC3^G31A^ hCerOs show reduced event frequency relative to EXOSC3^WT^, whereas mean inter-spike interval is not significantly different among genotypes. Data are ± SEM; each point is an independent organoid. Statistical significance was determined using one-way ANOVA with Tukey’s multiple comparison test; ns, not significant; ***p<0.0001. **(C)** Pairwise Spearman correlation matrix of spontaneous calcium activity among active ROIs from EXOSC3^WT^, EXOSC3^G31A^, and EXOSC3^G191C^ hCerOs. ROIs are grouped by genotype and hierarchically clustered within each group; red indicates positive (red) and blue (blue) indicates negative correlations. Mean pairwise Spearman correlation coefficients (ρ) are indicated for each genotype. **(D)** Organoid-level analysis of network synchrony. Cumulative distribution functions show mean pairwise Spearman correlation coefficients calculated independently for each organoid (n = 15 organoids per genotype). Shaded regions indicate pointwise 95% confidence bands obtained by organoid-level bootstrap resampling. Median organoid-level correlations and adjusted pairwise P values are indicated in the panel. **(E)** Representative spatial projection of calcium activity in EXOSC3^WT^, EXOSC3^G31A^, and EXOSC3^G191C^ hCerOs. Isochrone maps show the relative arrival time of calcium activity wave-responsive ROIs within a common 512 × 512-pixel imaging field. Color indicates arrival time relative to the first responding ROI, and white vectors indicate the direction of increasing arrival time. Maps are representative examples selected to illustrate differences in the spatiotemporal organization of calcium-wave propagation and are not used for genotype-level statistics. **(F)** Temporal coupling of individual calcium events to population-level activity. The population calcium signal was band-pass filtered at 0.01-0.20 Hz, and individual calcium events were assigned to the corresponding phase of the population activity cycle. Polar histograms show the cohort-balanced distribution of calcium events by population phase for EXOC3^WT^, EXOSC3^G31A^, and EXOSC3^G191C^ hCerOs (n = 15 organoids per genotype). Arrows indicate the circular mean vector, with direction representing preferred phase and vector length reflecting phase concentration. EXOSC3^WT^ events show greater concentration near the population-activity peak, whereas events in EXOSC3 mutant hCerOs are more broadly distributed throughout the population activity cycle.

We next asked whether *EXOSC3* mutations affect the coordination of activity within developing neuronal networks. Pairwise correlations among active regions of interest reveal substantially greater coordinated activity in EXOSC3^WT^ organoids than in either mutant genotype (**Fig. 4C**). Mean pairwise Spearman correlations decreased from ρ = 0.353 in EXOSC3^WT^ to ρ = 0.132 in EXOSC3^G191C^ and ρ = 0.088 in EXOSC3^G31A^ organoids. To determine whether this difference is reproducible among independent organoids rather than driven by individual cellular measurements, we calculated mean pairwise correlations at the organoid level. The distribution of organoid-level correlations shifted toward lower values in both mutant genotypes, with EXOSC3^G31A^ showing the largest reduction (n = 15 independent organoids per genotype; **Fig. 4D**). Complementary organoid-level analyses further reveal reduced recruitment and coordination of network activity in mutant hCerOs, including decreases in the fraction of event-positive cells, global synchronization, strong-edge coupling, and connectivity matching (**Fig. S5B**). Together, these independent measures support a broader disruption of network recruitment and coordination following EXOSC3 dysfunction.

We further examined the spatial organization of calcium activity by mapping the relative timing of cellular recruitment during representative propagation events (**Fig. 4E**). EXOSC3^WT^ activity propagates rapidly through the image field, with a representative event spanning 4.2s and an apparent propagation speed of 92.8μm/s. In comparison, representative EXOSC3^G191C^ and EXOSC3^G31A^ events exhibit progressively longer recruitment intervals and slower apparent propagations, spanning 5.9 and 14.4s with fitted speeds of 43.7μm/s and 19.4μm/s, respectively (**Fig. 4E**). Although these representative events illustrate differences in the spatiotemporal organization of network activity, they were selected to visualize fast, intermediate, and slow propagation patterns and therefore do not establish genotype-level differences in propagation speed.

Finally, given that neuronal activity in human brain organoids can become phase-coupled to population oscillations^71^, we asked whether individual calcium events are temporally organized relative to population-level activity. We therefore determined the phase of individual calcium events relative to the population activity cycle in each genotype (**Fig. 4F**). In EXOSC3^WT^ organoids, events preferentially occurred near the peak of population activity (mean angle, -13°; mean resultant length, 0.17). By contrast, calcium events in EXOSC3^G191C^ and EXOSC3^G31A^ organoids are more broadly distributed throughout the population cycle, with reduced phase concentration (20°, 0.07 and 57°, 0.09, respectively). Thus, cellular activity in EXSOC3^WT^ networks shows greater temporal coupling to the population rhythm, whereas this organization is reduced in both mutant genotypes.

Together, these analyses reveal that EXOSC3 dysfunction affects multiple levels of neuronal network organization. Mutant hCerOs generate spontaneous neuronal activity, but exhibit reduced cellular recruitment and network synchrony, altered spatiotemporal propagation, and diminished coupling of individual cellular events to population-level activity. These findings indicate that RNA exosome dysfunction does not prevent the acquisition of neuronal activity but disrupts its organization into coordinated network behavior.

### RNA exosome dysfunction alters the temporal coordination of lineage-specific developmental programs

To determine how EXOSC3 dysfunction alters transcriptomic landscapes during cerebellar differentiation, we modeled genotype- and maturation-associated changes in gene exwpression within individual cerebellar lineages. Mixed-effects analysis reveals that the dominant transcriptomic signature was associated with developmental progression from D35 to D60, with 4,651 transcripts classified as time-dependent and maturation-driven (**Fig. 5A)**. By comparison, fewer transcripts are associated predominantly with genotype-specific differences at D35 or genotype-by-maturation interactions. EXOSC3^G31A^ accounted for substantially more associated transcripts than EXOSC3^G191C^, consistent with a graded effect of the two PCH1b-associated variants unevenly among cerebellar populations, with Purkinje cells and PAX2+ interneuron progenitors (PIPs) contributing prominently to the affected transcript sets, followed by glia, rhombic lip, granule cell precursors, and granule neuron populations (**Fig. 5A**).

**Figure 5.**
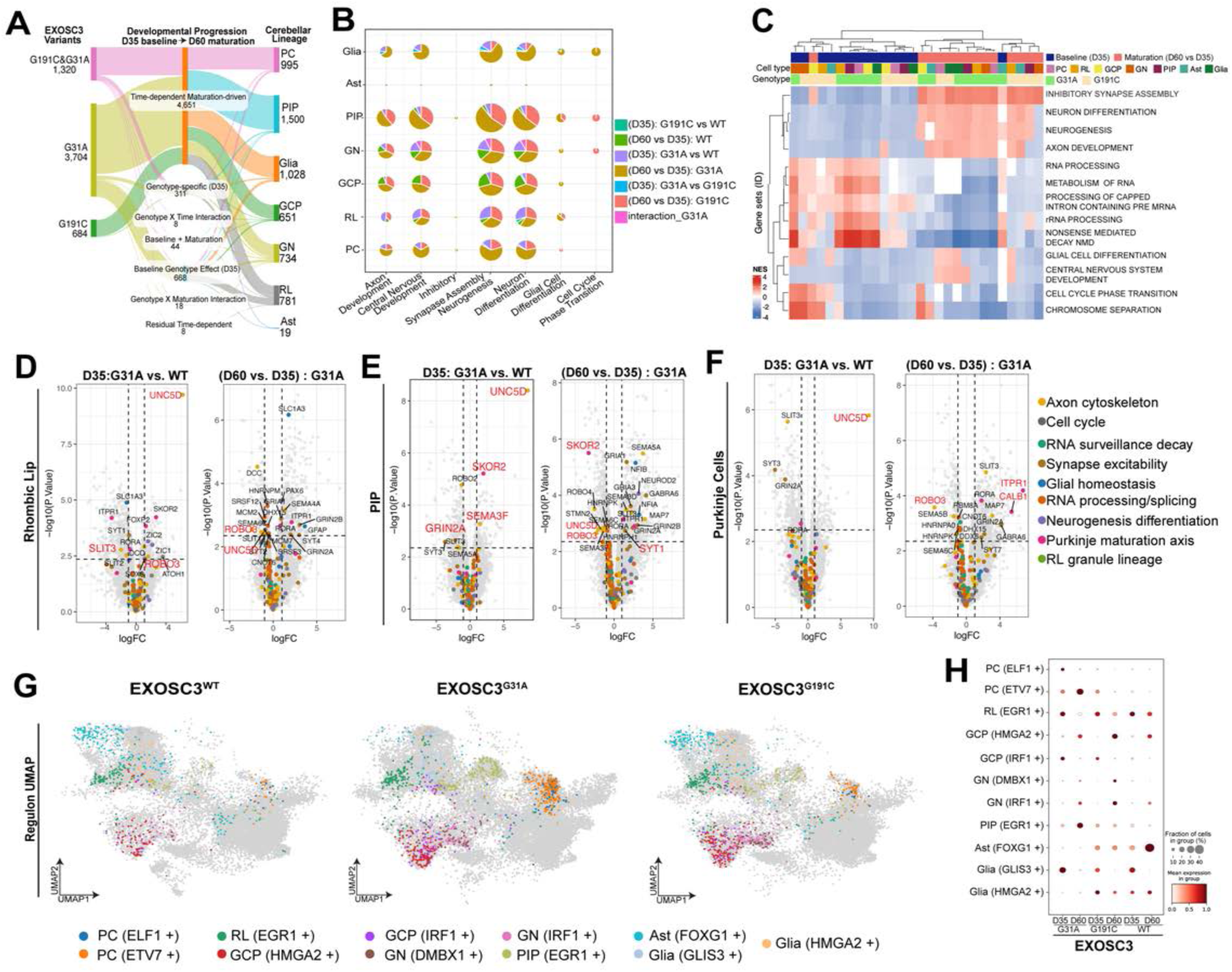
RNA exosome dysfunction alters the temporal coordination of lineage-specific developmental programs. **(A)** Sankey diagram summarizing lineage-resolved transcriptomic changes associated with EXOSC3^WT^, EXOSC3^G31A^, and EXOSC3^G191C^ during cerebellar development. Transcripts are classified according to baseline genotype effects at D35, time-dependent changes during D35-to-D60 maturation, and genotype-by-time interactions, and mapped to the indicated cerebellar lineages. Flow width represents the number of associated transcripts. Maturation-dependent changes comprise the largest transcriptomic class. **(B)** Lineage-resolved Gene Ontology (GO) enrichment analysis showing biological processes associated with genotype and developmental-time comparisons. Rows indicate cerebellar cell populations and columns indicate selected biological processes. Pie chart segments indicate the comparison contributing to each enrichment, as defined in the key; pie size is proportional to the number of differentially expressed transcripts contributing to each term. **(C)** Heatmap of gene set enrichment analysis (GSEA) of developmental and genotype-associated transcriptomes. Rows indicate selected biological processes, including neuronal differentiation, neurogenesis, axon development, synaptic organization, RNA processing, glial differentiation, and cell cycle regulation. Columns represent individual cell type, genotype, and developmental stage comparisons, as indicated by the annotation bars above the heatmap. Color represents normalized enrichment score (NES), with red indicating positive and blue indicating negative enrichment. **(D) – (F)** Lineage-resolved differential gene expression analysis in rhombic lip (D), PAX2+ interneuron progenitor (PIP; E), and Purkinje cell (F). For each lineage, the left volcano plot shows EXOSC3^G31A^ versus EXOSC3^WT^ at D35, and the right shows maturation-associated changes in EXOSC3^G31A^ between D60 and D35. Transcripts are plotted according to log_2_ fold change and -log_10_(p-value). Selected transcripts are labeled, and colored points indicate functional categories defined in the key: axon/cytoskeletal regulation, cell cycle, RNA surveillance/decay, synaptic/excitability programs, glial homeostasis, RNA processing/splicing, neurogenesis /differentiation, Purkinje maturation, and rhombic lip/granule-lineage programs. **(G)** UMAP projections showing representative lineage-associated regulon identities in EXOSC3^WT^, EXOSC3^G31A^, and EXOSC3^G191C^ hCerOs. Colors denote the indicated regulon-associated cell populations, including Purkinje cell, rhombic lip, granule cell precursor, granule neuron, PIP, astrocytic, and glial populations; cells not assigned to the displayed regulons are shown in gray. **(H)** Dot plot showing representative regulon activity among cell populations, genotypes, and developmental stages. Rows indicate lineage-associated regulons and columns indicate genotype and developmental timepoint (D35 and D60). Dot size represents the fraction of cells associated with each regulon, and color represents normalized regulon activity.

We next asked which biological processes are associated with these transcriptomic changes. Lineage-resolved enrichment analysis reveals prominent contributions from neuronal differentiation and neurogenesis, axon development, central nervous system development, and synaptic programs and cell cycle-associated pathways (**Fig. 5B**). Importantly, the relative contributions of baseline genotype and maturation-associated comparisons differ among cell types and biological processes. Gene set enrichment analysis similarly separated baseline and maturation-associated transcriptomic states, identifying coordinated changes in neuronal differentiation, neurogenesis, axon development, inhibitory synapse assembly, RNA processing, nonsense - mediated decay, cell cycle progression, and glial differentiation programs (**Fig. 5C**). Notably, RNA processing and surveillance-associated pathways were more prominent at D35, whereas neuronal differentiation and maturation-associated programs were more prominent at D60.

Lineage-resolved differential expression further localized these effects (**Fig. 5D-F, Fig. S6A-F**). In rhombic lip cells, EXOSC3^G31A^ organoids exhibit substantial transcriptomic dysregulation at D35, including altered abundance of transcripts associated with axon guidance, neuronal differentiation, and rhombic lip granule-lineage development (**Fig. 5D**). *UNC5D* is among the most strongly increased transcripts at D35, accompanied by changes in developmental regulators and guidance-associated transcripts including *SKOR2*, *ROBO3*, and *SLIT3*. Comparing D60 with D35 in EXOSC3^G31A^ hCerOs shows that many of these transcripts remain developmentally responsive, with several changing in abundance as differentiation proceeds (**Fig. 5D**). These observations argue against a uniform failure to respond in early transcriptomic states and instead indicate altered timing and magnitude of developmental gene expression programs.

PAX2+ interneuron progenitors similarly display genotype-dependent transcriptomic differences during the neurogenesis window **(Fig. 5E**). At D35, EXOSC3^G31A^ hCerOs show altered abundance of neuronal differentiation-associated transcripts including *SKOR2*, *SEMA3F*, *GRIN2A*, *ROBO2*, and *UNC5D*. Notably, *SKOR2*, a transcriptional regulator implicated in Purkinje cell specification and suppression of interneuron identity, exhibits increased in EXOSC3^G31A^ Purkinje lineage intermediates (PIPs) at D35 but decreased during subsequent maturation. This transient detection of lineage-associated transcriptional program outside its predominant developmental context suggests reduced temporal or lineage restriction of gene expression rather than irreversible acquisition of an alternative cell fate.

Purkinje cells exhibit a distinct transcriptomic response (**Fig. 5F**). Although baseline differences at D35 are comparatively restricted, maturation is accompanied by substantial remodeling of Purkinje cell gene expression in EXOSC3^G31A^ organoids. Transcripts associated with Purkinje cell maturation and neuronal function, including *RORA*, *CALB1*, *ITPR1*, and *GRIN2A*, increase between D35 and D60, while axon guidance-associated transcripts are differentially regulated, including increased *SLIT2* and decreased *ROBO3* (**Fig. 5F**). Thus, despite EXOSC3 dysfunction, Purkinje cells continued to express maturation-associated transcripts but followed altered transcriptomic trajectories. These findings are consistent with the cellular analyses showing that mutant Purkinje-lineage cells are generated but progress through developmental states abnormally rather than undergoing a complete developmental arrest.

To independently validate differential expression patterns identified by scRNA-seq, we performed RT-qPCR on D35 EXOSC3^WT^ and *EXOSC3* mutant hCerOs. Because RT-qPCR measures transcript abundance in bulk organoid samples, we selected candidates exhibiting consistent differential expression in multiple cell populations rather than changes restricted to individual lineages. *UNC5D* exhibits increased abundance in 20 of 21 defined clusters, *GPC5* exhibits increased abundance in 14 of 21 defined clusters, whereas *ZNF667* and *ZNF661-AS1* exhibit decreased abundance in 17 and 14 clusters, respectively, in D35 EXOSC3^G31A^ relative to EXOSC3^WT^ hCerOs. RT-qPCR confirms increased *GPC5* expression and decreased *ZNF667* and *ZNF667-AS1* in EXOSC3^G31A^ hCerOs, consistent with scRNA-seq findings (**Fig. S6G)**. These results provide independent validation of selected transcriptomic changes associated with EXOSC3 dysfunction.

We next examined whether these transcriptomic effects correspond to changes in cell type-associated regulatory states. Projection of representative regulon assignments onto the integrated hCerO landscape reveals distinct disruptions of regulatory programs associated with Purkinje cells, rhombic lip cells, GCPs, granule neurons, PIPs, astrocytes, and glia (**Fig. 5G**). The relative representation and distribution of these regulon-associated states differ among EXOSC3^WT^, EXOSC3^G31A^, and EXOSC3^G191C^ organoids, further indicating that RNA exosome dysfunction alters developmental regulatory states in a lineage-dependent manner. Examination of representative regulation at D35 and D60 similarly reveals genotype- and developmental stage-dependent differences among neuronal and glial populations (**Fig. 5H**).

Together, these analyses show that EXOSC3 dysfunction does not simply maintain an early transcriptomic state or globally suppress differentiation. Instead, RNA exosome dysfunction alters timing, magnitude, and lineage specificity of gene expression programs as cerebellar cells progress through differentiation. The strongest transcriptomic effects associated with EXOSC3^G31A^ further parallel the graded cellular and functional phenotypes observed. These findings prompted us to ask whether particular lineage-defining regulatory programs are selectively vulnerable to impaired RNA surveillance, which we addressed through regulon-level analysis (**Fig. 6**).

**Figure 6.**
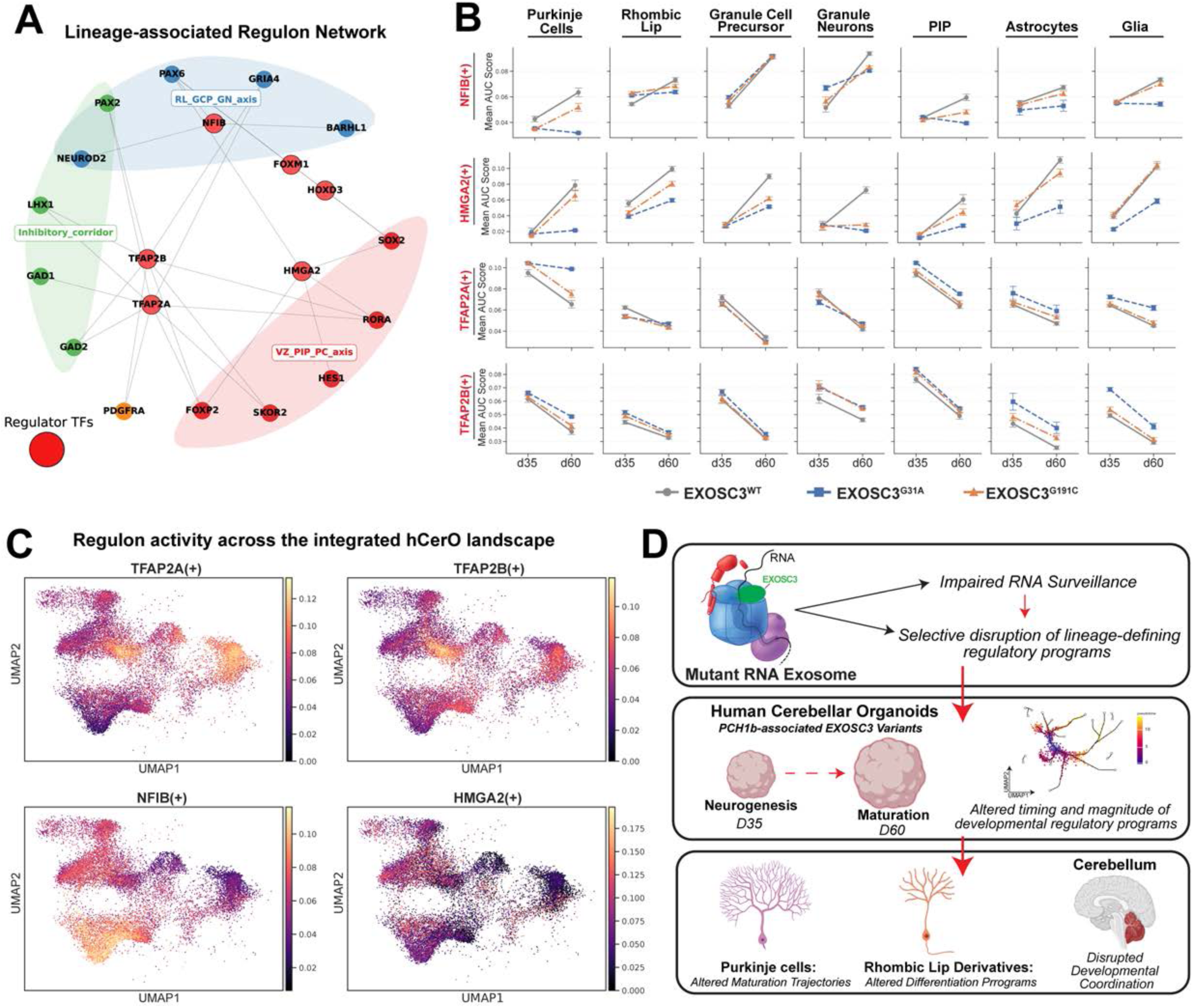
EXOSC3 dysfunction selectively alters lineage-associated regulatory programs during cerebellar maturation. **(A)** Lineage-associated gene regulatory network inferred from SCENIC regulon analysis. Nodes represent transcription factors (TFs) and predicted target genes, and edges indicate inferred regulatory relationships. Regulatory modules associated with the rhombic lip-granule cell precursor-granule neuron (RL-GCP-GN) axis, inhibitory/interneuron lineages, and ventricular zone-PIP-Purkinje cell (VZ-PIP-PC) axis are indicated by shaded regions. Red nodes oultlined in black denote representative regulatory TFs. **(B)** Lineage-resolved developmental trajectories of NFIB(+), HMGA2(+), TFAP2A(+), and TFAP2B(+) regulon activity in Purkinje cells, rhombic lip, granule cell precursors, granule neurons, PAX2+ interneuron progenitors (PIP), astrocytes, and glia. Mean regulon AUC scores are shown at D35 and D60 for EXOSC3^WT^, EXOSC3^G31A^, and EXOSC3^G191C^ hCerOs. Error bars indicate SEM. Global statistical analyses identified significant genotype-, developmental timepoint-, and cell type-associated interaction effects for regulons (**Table S2**). Genotype- and lineage-dependent differences in the direction and magnitude of developmental changes illustrate selective alteration of regulatory programs during maturation. **(C)** Regulon activity projected onto the integrated hCerO single-cell landscape. UMAPs show continuous AUC scores for TFAP2A(+), TFAP2B(+), NFIB(+), and HMGA2(+) regulons, illustrating the cellular distribution of each regulatory program within the integrated dataset. Color represents regulon AUC score as indicated by the corresponding scale. **(D)** Model summarizing the consequences of RNA exosome dysfunction during human cerebellar development. PCH1b-associated EXOSC3 variants impair RNA surveillance and are associated with selective disruption of lineage-associated regulatory programs. During the transition from neurogenesis (D35) to maturation (D60), these changes alter the timing and magnitude of developmental regulatory programs, contributing to altered Purkinje cell maturation trajectories, altered rhombic lip-derived differentiation programs, and disrupted coordination of cerebellar development.

### EXOSC3 dysfunction selectively alters lineage-associated regulatory programs during cerebellar maturation

To determine whether the lineage-specific developmental phenotypes associated with EXOSC3 dysfunction are accompanied by disruption of regulatory programs that define cerebellar cell states, we inferred transcription factor-associated regulons from the single-cell transcriptomic data. Regulon network analysis identifies modules that associate with major cerebellar developmental axes, including an RL-GCP-GN associated network containing NFIB and a VZ-PIP-PC associated network containing TFAP2A, TFAP2B, FOXP2, SKOR2, RORA, and HES1 (**Fig. 6A**). These findings identify candidate regulatory programs through which broadly impaired RNA surveillance could differentially affect individual cerebellar lineages.

We next examined regulon activity among EXOSC3^WT^, EXOSC3^G31A^, and EXOSC3^G191C^ hCerOs at D35 and D60 in Purkinje cells, rhombic lip cells, granule cell precursors, granule neurons, PAX2+ interneuron progenitors, astrocytes, and glia (**Fig. 6B, Fig. S7**). Regulon activity changes substantially during maturation in EXOSC3^WT^ organoids, but the magnitude and direction of these developmental trajectories differ following EXOSC3 disruption. NFIB-associated activity, for example, increases during EXOSC3^WT^ Purkinje cell maturation but fails to show a comparable increase in EXOSC3^G31A^ cells, whereas EXOSC3^G191C^ displays an intermediate trajectory. Similar genotype-dependent attenuation of developmental changes in NFIB regulon activity is observed in PIPs and glial populations. HMGA2-associated activity likewise increases between D35 and D60 in several EXOSC3^WT^ lineages, including Purkinje cells, rhombic lip cells, GCPs, PIPs, astrocytes, and glia, while these maturation-associated increases are attenuated to differing degrees in *EXOSC3* mutant cells (**Fig. 6B**). Notably, these effects are not uniform among the lineages: granule neurons exhibit a distinct HMGA2 trajectory, emphasizing the cell type-dependent consequences of EXOSC3 dysfunction.

This selectivity is also evident among related regulatory programs. TFAP2A- and TFAP2B-associated activities generally decrease during maturation, but their response to EXOSC3 disruption differs by cell type (**Fig. 6B**). In Purkinje cells, TFAP2A activity remains elevated in EXOSC3^G31A^ cells at D60 relative to the developmental decline observed in EXOSC3^WT^ and EXOSC3^G191C^ cells, whereas TFAP2B shows a comparatively similar maturation-associated decline among genotypes. Conversely, genotype-dependent differences in TFAP2B activity are more apparent in astrocytic and glial populations. Thus, EXOSC3 dysfunction does not produce a uniform shift in transcription factor-associated activity but instead alters specific regulatory trajectories within particular developmental contexts.

Projection of representative regulon activities onto the integrated hCerO embedding further demonstrates that TFAP2A, TFAP2B, NFIB, and HMGA2 activities are spatially restricted within the cellular landscape rather than uniformly distributed among cerebellar populations (**Fig. 6C**). Together with the lineage-resolved developmental trajectories, these patterns indicate that EXOSC3 dysfunction affects both developmental stage and the regulatory architecture of individual cell types.

Collectively, these analyses reveal that ubiquitous disruption of RNA exosome function is associated with selective dysregulation of lineage-defining regulatory programs rather than a uniform collapse of transcriptional regulation (**Fig. 6D**). *EXOSC*3 mutations alter the timing and magnitude of regulon activity during cerebellar maturation, with prominent effects in Purkinje cells and additional lineage-specific consequences in rhombic lip-derived and glial populations. These findings support a model in which RNA exosome-mediated surveillance contributes to the temporal coordination of developmental regulatory programs, providing a potential mechanism by which disruption of a ubiquitously required RNA processing and decay complex produces selective developmental vulnerability in the human cerebellum.

## Discussion

Cellular differentiation requires precise coordination of gene expression programs over developmental time^17, 31, 35, 36^, yet how co- and post-transcriptional RNA surveillance contributes to this process remains poorly understood. Here, we identify RNA exosome-mediated surveillance as an important contributor to the temporal coordination of human cerebellar development. Major cerebellar lineages remain specified and continue to progress through developmental time following EXOSC3 disruption, but their developmental trajectories, relative abundance, spatial organization, and maturation-associated transcriptional programs are altered. The consequences of RNA exosome dysfunction are particularly pronounced in Purkinje cells and rhombic lip-derived populations, with accompanying disruptions in lineage-associated regulatory programs and impaired coordination of neuronal network activity. Together, these findings provide a framework in which dysfunction of a ubiquitously expressed RNA surveillance complex produces selective developmental vulnerability in the human nervous system and support an early neurodevelopmental component of PCH1b pathogenesis.

Cell fate transitions are often viewed through a transcription-centric lens; however, RNA-level regulation is increasingly recognized as an important determinant of developmental progression, particularly in the central nervous system (CNS)^32, 72, 73^. Neuronal development requires tightly coordinated RNA processing^36, 53, 74^, localization^75–77^, and turnover^78^ as cells transition between progenitor and differentiated identities. While RNA processing and localization have been extensively studied in CNS development and disease^37, 38, 79, 80^, the contribution of RNA decay, particularly RNA exosome-dependent surveillance^17, 27, 28^, remains comparatively underexplored. Other RNA surveillance pathways provide precedent for such developmental functions; for example, disruption of nonsense-mediated decay (NMD) permits neural stem cell proliferation but impairs subsequent neuronal differentiation^81^. These observations suggest that RNA surveillance can influence developmental progression beyond its canonical role in eliminating aberrant RNAs.

The RNA exosome is a conserved ribonuclease complex responsible for degrading diverse coding and non-coding RNAs^16, 17, 34^. Although frequently considered a general RNA quality control machine, our findings support a broader role for the RNA exosome during developmental transcriptome remodeling. Importantly, the effects of EXOSC3 dysfunction cannot be explained by a uniform failure to eliminate early developmental transcripts. Instead, lineage-resolved analyses reveal distinct temporal behaviors among affected genes. Transcripts including ^82^, *ROBO3*, and *SKOR2* are elevated in specific populations at D35 but remain developmentally responsive, decreasing during subsequent maturation, whereas Purkinje cells activate maturation-associated genes including *RORA*, *CALB1*, *ITPR1*, and *GRIN2A*. Thus, developmental programs continue to change in mutant cells, but their timing, magnitude, and cellular context differ from wildtype. These findings suggest that RNA exosome function contributes to the coordinated remodeling of gene expression programs rather than acting simply as an on/off mechanism for developmental transcript clearance.

This lineage specificity provides insight into a central feature of RNA exosomopathies: mutations affecting ubiquitous RNA processing machinery produce selective neurological phenotypes^8, 42, 83^. Consistent with this principle, previous work in erythroid progenitors demonstrated that disruption of RNA exosome function produces pronounced lineage-specific defects despite relatively restricted changes in steady-state RNA abundance^84^. *EXOSC3* mutant hCerOs generate the major cerebellar populations, arguing against a generalized failure of lineage specification, but show lineage-dependent alterations in developmental trajectories and gene expression programs. Purkinje cells, rhombic lip-derived populations, and interneuron progenitors are particularly affected, whereas several glial populations show comparatively limited transcriptomic changes. Consistent with this selectivity, regulon analysis reveals genotype- and developmental stage-dependent alterations in NFIB(+), HMGA2(+), TFAP2A(+), and TFAP2B(+) activity rather than uniform disruption of transcriptional regulation. These findings suggest that the consequences of impaired RNA surveillance depend on the regulatory programs being executed within particular developmental contexts, providing a principal explanation for selective neuronal vulnerability despite ubiquitous RNA exosome expression.

In addition to this lineage specificity, our findings reveal a temporal distinction in the molecular consequences of RNA exosome dysfunction. RNA processing and surveillance-associated pathways are more prominent among transcriptomic alterations at D35, whereas neuronal differentiation and maturation programs become more prominent at D60. This temporal pattern is consistent with a model in which early perturbations in RNA homeostasis precede and potentially contribute to subsequent defects in neuronal lineage progression. Although these findings do not establish causality, they suggest that early disruption of RNA regulatory programs may have lasting consequences for cerebellar development.

Altered developmental progression is accompanied by disrupted cerebellar tissue organization. *EXOSC3* mutant hCerOs generate BARHL1+ granule lineage cells and CALB1+ Purkinje cells but show increased ectopic BARHL1+ cells within or below the Purkinje cell layer. Because Purkinje cells regulate granule cell precursor proliferation and differentiation through pathways including Sonic hedgehog (Shh)^85–87^, the coupled abnormalities in Purkinje and rhombic lip-derived populations raise the possibility that altered maturation of one lineage influences development of another. Determining whether altered Purkinje-derived signaling contributes to granule lineage phenotypes will be an important future direction.

These developmental abnormalities extend to neuronal network function. All genotypes acquire spontaneous calcium activity, but EXOSC3^G31A^ hCerOs show reduced event frequency, and both mutant genotypes exhibit reduced coordination of spontaneous activity. Mutant networks show reduced activity, indicating disruption of both cell-to-cell and cell-to-population temporal organization. Thus, EXOSC3 dysfunction does not prevent neurons from becoming active but compromises the organization of that activity into coordinated network behavior.

Together, these findings connect RNA regulation to developmental progression, tissue organization, and neuronal function. Rather than producing uniform developmental arrest, EXOSC3 dysfunction allows differentiation to proceed while altering the timing, magnitude, and lineage specificity of developmental programs. This model helps reconcile the ubiquitous requirement for RNA exosome function with selective vulnerability in RNA exosomopathies: the consequences of impaired RNA surveillance may depend on the developmental programs and RNA processing demands encountered by individual cell populations.

Our findings provide mechanistic insight into how RNA exosome dysfunction alters developmental progression, while the molecular determinants underlying transcript and lineage selectivity remain to be defined. Transcriptomic changes provide a readout of altered RNA exosome function but cannot distinguish direct effects on RNA turnover from secondary changes in transcription or developmental regulation. Thus, altered transcript abundance may reflect changes in RNA stability, transcription, or both. Defining direct RNA exosome substrates and their decay kinetics will require complementary approaches that directly measure RNA synthesis, turnover, and RNA exosome-substrate interactions. Critically, the emergence of molecular, cellular, and tissue-level abnormalities during neurogenesis and early maturation in hCerOs modeling PCH1b-associated EXOSC3 variants supports an early neurodevelopmental component of PCH1b. More broadly, our findings establish RNA surveillance as an important contributor to the temporal coordination of developmental gene expression programs, providing a framework for how disruption of ubiquitous RNA processing and decay machinery can produce selective vulnerability during human brain development.

### Limitations of the Study

Although our findings establish a role for RNA exosome function in coordinating cerebellar developmental programs, several limitations remain. Our transcriptomic analyses cannot distinguish direct effects on RNA turnover from secondary changes in transcription or developmental regulation. Furthermore, poly(A)-based scRNA-seq does not comprehensively capture all RNA exosome substrates, including non-polyadenylated transcripts. Moreover, although cerebellar organoids recapitulate key features of early cerebellar development, they do not fully reproduce the cellular complexity and maturation of the developing human cerebellum.

## KEY RESOURCES TABLE

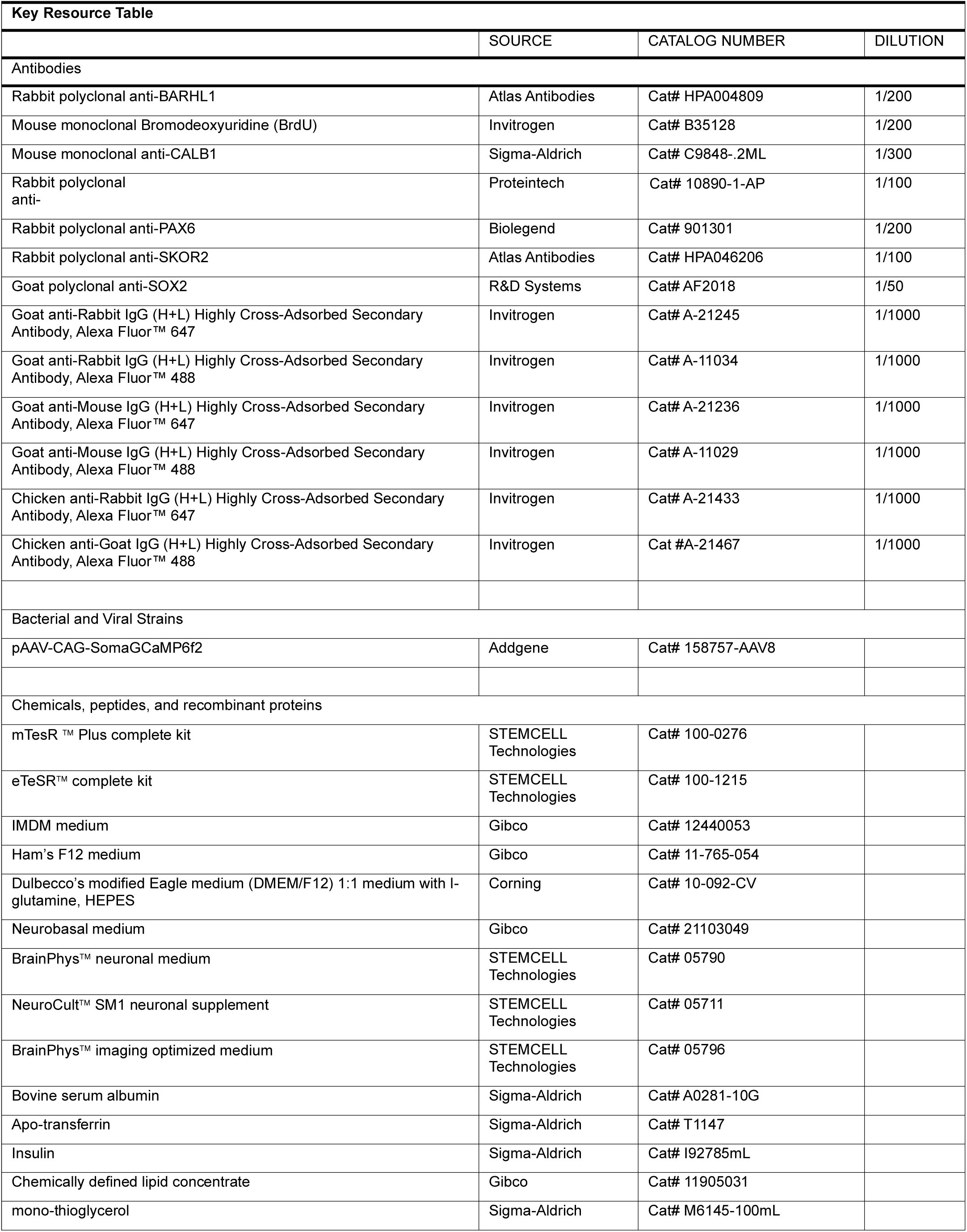
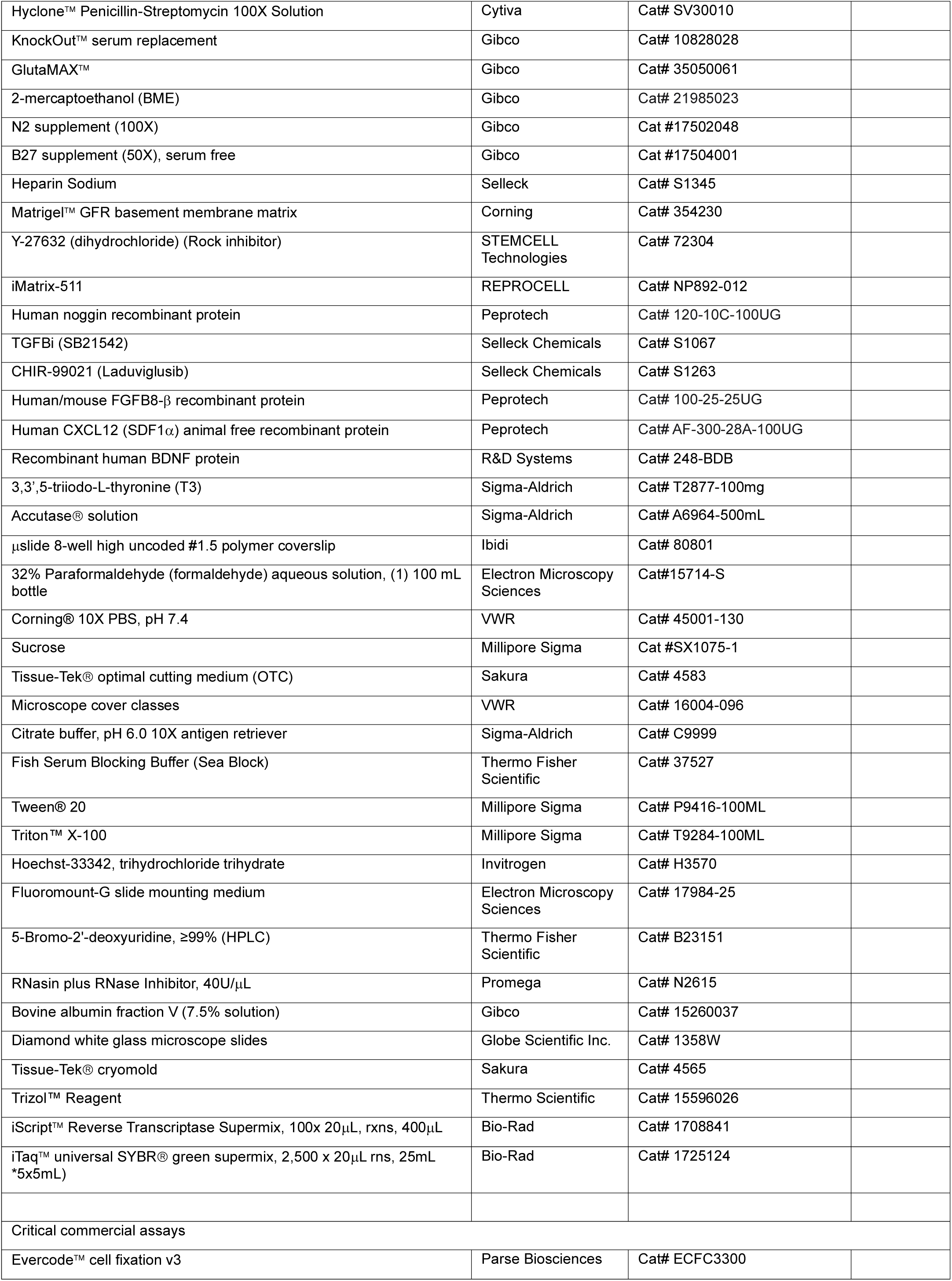
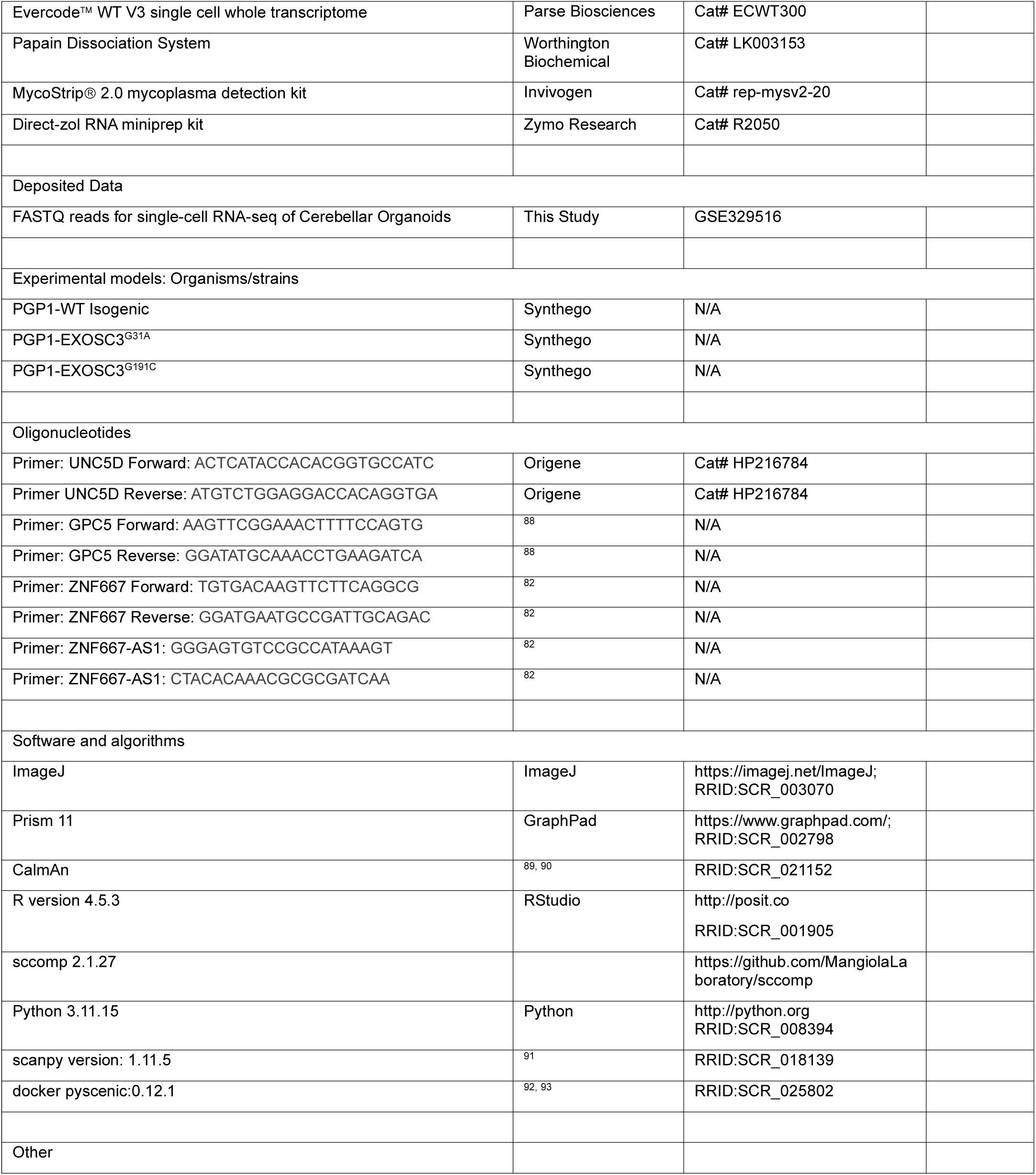

## EXPERIMENTAL MODEL AND STUDY PARTICIPANT DETAILS

### Generation of CRISPR-edited stem cell lines

The PGP1 (Personal Genome Project 1) human iPSC line was edited using CRISPR/Cas9 to generate EXOSC3^G31A^ and EXOSC3^G191C^ PCH1b mutant lines (Synthego). An isogenic control line was generated in parallel using the same editing workflow with a non-targeting (blank) CRISPR cassette. Two independently derived clonal populations were established for each genotype. Genome editing was confirmed by Sanger sequencing, and clones were expanded independently before differentiation (**Fig. S3**). All experiments incorporated two independent clones per genotype and independent organoid differentiations as described below.

Cells were cultured in mTeSR Plus or eTeSR medium (STEMCELL Technologies) supplemented with 100 U/mL penicillin and 100 μg/mL streptomycin (Cytiva) at 37°C with 5% CO2. Passages were completed utilizing Accutase (Sigma-Aldrich) with media supplemented with 0.25 μg/cm^2^ iMatrix (REPROCELL) and 10 μM ROCK inhibitor (STEMCELL Technologies). Cultures were maintained below passage 65 and routinely tested negative for mycoplasma (Invivogen).

### Experimental design and definition of biological replicates

Independent organoid differentiations were initiated from separately maintained hiPSC cultures and were considered independent biological experiments. Within each differentiation, individual organoids were generated in separate wells and subsequently maintained as described below. For imaging experiments, multiple sections and/or regions of interest obtained from the same organoid were treated as within-organoid measurements and were not considered independent biological replicates. We summarized measurements at the organoid level, where appropriate, before statistical comparisons between genotypes. The number of independent differentiations, organoids, and within-organoid measurements used for each experiment is reported in the corresponding figure legend.

For single-cell RNA sequencing, each biological replicate consisted of an independently collected pool of 8-12 organoids. Two biological replicate pools were generated for each genotype at each developmental time point and were processed independently through dissociation, fixation, library preparation, and sequencing. Individual cells represent observations nested within biological replicate samples and were not considered independent biological replicates.

## METHOD DETAILS

### Cerebellar Organoid Generation

Human cerebellar organoids were generated as described in Atamian *et al.* 2025 *Nature Protocols*^58^ with minor modifications. Human (PGP1) iPSCs were cultured to approximately 80% confluency, dissociated into single cells, and seeded at 6,000 cells per well in ultra-low attachment 96-well V-bottom plates. Cells were aggregated in gfCDM supplemented with 10μm TGF-β inhibitor (Selleck Chemicals), 1.7μm CHIR (Selleck Chemicals), 50ng/ml Noggin (Peprotech), and 20μm ROCK inhibitor to promote neuronal induction. 48 hours after seeding, we removed half the media and replenished it with Noggin at an increased concentration of 150 ng/ml.

During early differentiation, organoids were patterned by supplementing with morphogens, including CHIR, Noggin, and FGF8β (Peprotech), to promote midbrain-hindbrain boundary identity. On day 4, a half media change was performed with fresh gfCDM supplemented with 10μm TGF-β inhibitor, 1.7μm CHIR, 100ng/ml Noggin, and 200ng/ml FGF8β. Starting on day 6, the organoid media was transitioned to CerDM1 with the same morphogen concentrations except FGF8β was decreased to 100ng/ml. On day 8, 80 μl of media was removed, and an additional 100 µl of CerDM1 supplemented with FGF8β to 400ng/ml, TGF-β, CHIR, and Noggin remained consistent. On day 10, 80 μl of media was removed, and an additional 100 μl of CerDM1 supplemented with 300ng/ml FGF8β, and TGF-β, CHIR, and Noggin remained consistent. On days 12 and 14, 70 µl was removed and replaced with 75 µl of CerDM1. On day 16, organoids were transferred to suspension culture and cultured in CerDM2 media. From day 30, organoids were maintained in maturation media (CerDM3) supplemented with 1% Matrigel (Corning). 100ng/ml SDF1α (Peprotech) and 0.5ng/ml T3 (Sigma-Aldrich) were added to support cerebellar lineage specification and tissue organization.

Media were exchanged every 3 days. Organoids were collected at D35, representing an active neurogenic stage, and D60, representing early neuronal maturation. For long-term functional experiments, organoids were maintained beyond D60 in maturation medium supplemented with 0.5ng/ml T3 and 14ng/ml BDNF (R&D Systems), with media exchanged every 3-4 days. Organoids for each genotype were generated using the same differentiation protocol and experimental schedule.

### Cryosectioning and Immunofluorescence

Organoids were collected from culture dishes and fixed with 4% paraformaldehyde (Electron Microscopy Sciences) for 30 min at room temperature, followed by three washes in PBS (VWR). Samples were cryoprotected in 30% sucrose (Millipore Sigma) in PBS overnight, embedded in Tissue-Tek O.C.T. compound (Sakura), and stored at -80°C. Organoids were cryosectioned at 14 μm onto glass slides (Globe Scientific).

Sections were permeabilized with 0.3% Triton-X (Millipore Sigma) PBS and washed three times, followed by antigen retrieval in citrate buffer (Sigma-Aldrich) using a pressure cooker for 12 min. After washing, sections were blocked for 1 hour at room temperature in PBS containing 0.3% Triton-X PBS and 1.5% Sea Block (Thermo Fisher Scientific). Primary antibodies were diluted in the antibody buffer (1.5% Sea Block (Thermo Fisher Scientific), 0.1% Tween-20 (Millipore-Sigma) in PBS) and incubated overnight at 4°C.

Following three washes in 0.1% Tween-20 in PBS, sections were incubated in secondary antibody diluted in antibody buffer. Slides were washed three times, counterstained with Hoechst (Invitrogen) for 30 min, washed again, and mounted using Fluoromount-G (Electron Microscopy Services) with glass coverslips (VWR). Primary and secondary dilutions are specified in the Key Resource Table.

### BrdU proliferation assay

For proliferation analysis, 7-9 organoids were transferred to ultra-low attachment 6-well plates and incubated in CerDM3 supplemented with 100μM bromodeoxyuridine (BrdU) (Thermo Fisher Scientific). Organoids were labeled for 2 hours at 37 °C with 5% CO_2_ on an orbital shaker (70rpm). Followed by three washes in CerDM3. Organoids were then cultured in CerDM3 supplemented with 1% Matrigel, 100ng/mL SDF1α, and 0.5ng/mL T3 for 24 hr. Samples were fixed in 4% paraformaldehyde, cryoprotected in 30% sucrose, embedded, and cryosectioned as described above. BrdU incorporation was assessed by immunofluorescence.

### Microscopy and image analysis

Organoids were imaged using Leica DMi8, Leica Stellaris, or MICA imaging systems using identical acquisition settings within each experiment marker combination. Image analysis was performed using ImageJ/Fiji (NIH). For each organoid, sections spaced approximately 140 μm apart were imaged and analyzed. Measurements from individual sections were averaged to generate a single biological replicate per organoid. Multiple regions obtained from the same organoid were considered within-organoid statistical comparisons rather than independent replicates. Where genotype-level statistical comparisons were performed, measurements from multiple regions were averaged to generate a single value per organoid. Independent organoids served as biological replicates. The number of organoids, independent differentiations, sections, and regions analyzed for each experiment is reported in the corresponding figure legend.

### Dissociation of cerebellar organoids and single-cell RNA sequencing

Organoid dissociation was adapted from Atamian *et al.*^58^ and the Worthington Papain dissociation protocol. Briefly. 8-12 organoids were collected and mechanically minced, followed by enzymatic dissociation using papain supplemented with DNase I and RNase inhibitor (Promega). Samples were incubated at 37°C with agitation and triturated to achieve a single-cell suspension.

Cells were purified using an ovomucoid inhibitor solution and centrifugation (300 x g, 7 min), then resuspended in PBS containing 0.04% bovine serum albumin (Gibco) and filtered through a 40μm strainer (Corning). Cell counts were determined, and 150,000 and 500,000 cells per clonal replicate per genotype were fixed and permeabilized using the Parse Biosciences Cell Fixation and Permeabilization Kit according to the manufacturer’s instructions. Fixed cells were stored at -80°C until processing.

Single-cell libraries were generated using the Parse Biosciences Evercode WT v3 kit. Approximately 8,300 cells per biological replicate (n=2 per genotype per timepoint) were barcoded and sequenced on a NovaSeq X Plus platform.

### Single-cell RNA-sequencing processing and analysis

Single-cell RNA-seq was performed using two independently derived hiPSC clones per genotype at D35 and D60. For each clone and developmental timepoint, 8-12 cerebellar organoids were pooled to generate one biological replicate. Thus, each genotype and timepoint was represented by two biological replicates derived from distinct hiPSC clones. Samples were processed independently before integration for downstream analysis. Biological replicate identity was retained in the metadata throughout processing and downstream statistical analyses. Replicates were integrated for visualization and cell type annotation where indicated.

### Quality Control, Normalization, and Dimensionality Reduction

Single-cell RNA-seq data were processed using Scanpy^91^. Cells were filtered based on standard quality control metrics, including total UMI counts, number of detected genes, and mitochondrial transcript percentage. All samples showed high cell retention rates following filtering (97-99%; **Table S1**), indicating robust data quality and consistent capture across genotypes and developmental timepoints. Doublets were identified and removed using Scrublet with an expected doublet rate of 5%. Expression matrices were normalized and log-transformed. To mitigate technical and cell-cycle-associated variability, total counts per cell, mitochondrial transcript percentage, and cell-cycle scores (S and G2/M phases) were regressed out. Principal component analysis (PCA) was performed on the normalized matrix. Neighborhood graphs were constructed using the top principal components, and UMAP embeddings were generated to visualize cellular heterogeneity across all genotypes and time points. Potential batch effects were assessed via linear regression to estimate the contribution of sample identity to variance across all principal components; because batch effects were minimal, no additional correction (e.g., Harmony) was applied.

### Clustering and Cell Type Annotation

Cells were clustered using the Leiden algorithm (resolution = 1.0) to identify transcriptionally distinct populations. Cell identities were assigned by mapping query datasets to a human fetal cerebellum reference dataset using Scanpy’s ingest function. To reduce bias from highly abundant populations, an equal sampling strategy of 500 cells per cluster was performed prior to reference mapping.

### Trajectory Inference and Graph Abstraction

Trajectory analysis was performed using Scanpy on cells restricted to highly variable genes using log-transformed expression values. Quality control filtering retained cells with >200 and < 6,000 detected genes and <10% mitochondrial transcripts. Total counts and detected gene numbers were computed per cell, with metadata parsed from sample identifiers. Following PCA and k-nearest neighbor graph construction, clustering was performed using the Leiden algorithm (resolution = 0.5).

To model global connectivity between clusters, partition-based graph abstraction (PAGA) was applied using Leiden clusters as groups, with edges exhibiting connectivity < 0.05 excluded from visualization. A force-directed layout initialized from the PAGA topology was used to generate trajectory-informed embeddings. Cells were ordered along pseudotime based on the trajectory embedding. For centroid-based trajectory abstraction, cells were projected into a low-dimensional embedding and partitioned into coarse states using k-means clustering (k = 10, seed = 0. Cluster centroids defined graph nodes, and a minimum spanning tree (MST) based on Euclidean distances between centroids was constructed to represent global topology, with edges oriented from younger to older clusters based on mean cell age.

### Gene Regulatory Network and Regulon Analysis (pySCENIC)

Gene regulatory network analysis was executed using the SCENIC/pySCENIC workflow^92, 93^. Raw scRNA-seq counts were subset to selected cerebellar cell types (PC, Ast, PIP, RL, Glia, GCP, GN), with each cell type downsampled to a maximum of 500 cells to prevent overrepresentation. Genes detected in fewer than 10 cells were excluded, and the top 2,000 highly variable genes were retained. Transcription factors within this set and curated organoid-relevant target genes were used for downstream analysis, removing cells with zero total counts post-filtering.

Candidate transcription factor (TF)-target modules were refined using cisTarget motif enrichment, retaining motifs with normalized enrichment scores (NES) > 3.0 within ±10kb of transcription start sites. Directed TF-target networks were constructed with TFs as source nodes and target genes as downstream nodes, weighted by inferred regulatory strength. Regulon activity was quantified using SCENIC AUC scores. Dominant regulons were assigned based on maximum AUC, and activity was binarized using a 90th percentile threshold (excluding populations with <100 cells).

### Differential Composition and Temporal Mixed-Effects Modeling

#### Cellular Lineage Grouping

Fine-grained cell types were grouped into four major classes for lineage dynamics analysis: (1) Purkinje lineage (PC, PIP), (2) inhibitory/interneuron lineage (MLI, iCN, OPC), (3) glial lineage (Glia, Ast/Ependymal), and (4) Rhombic lip-derived lineage (RL, GCP, GN, eCN, UBC). Cells were binned along pseudotime, and cell counts per class were quantified within each bin to evaluate genotype-dependent differences.

#### Mixed-Effects Modeling

Cell type proportions were modeled using a Dirichlet-multinomial framework implemented in the sccomp R package^94^. Models incorporated genotype, time point, and their interaction as fixed effects, with the biological replicate pool treated as the experimental unit and random effect. Genotype-specific temporal effects were computed by combining baseline and interaction terms. Statistical significance was evaluated at FDR-adjusted p<0.05, and 95% CI were reported

### Calcium imaging and analysis

#### Data acquisition and sample preparation

Organoids were placed into an 8-well imaging plate (Ibidi) and transduced with pAAV-CAG-SomaGCaMP6f2 (Addgene, #158757-AAV8) at a final viral load of 1 x 10^11^ vg/mL in CerDM3. One day after infection, a full media change to BrainPhys Optimized Medium (STEMCELL Technologies) was conducted. After 7 days, the organoids were transferred to a temperature-controlled chamber supplied with 5% CO_2_ and imaged with a Leica SP-8 microscope using a multiphoton laser. The excitation wavelength was set to 920 nm, and the detector recorded emission spectra between 500 and 520 nm. Time-lapse recordings were acquired at 1.18 Hz, for 7.5 min, using a Fluotar VISIR 25x (0.95 NA) water objective, and exported as multi-frame TIFF files. Raw GFP imaging data were motion-corrected and analyzed using the constrained non-negative matrix factorization (CNMF) and the CaImAn computational toolbox as below and in^89, 90^. Spontaneous calcium activity was recorded from long-term hCerOs at approximately D150. Fifteen independent organoids were analyzed per genotype (n = 15 biological replicates per genotype).

#### Signal extraction and quality control

Calcium-imaging TIFF stacks were processed in CaImAn. Movies were motion-corrected using NoRMCorre, and neuronal regions of interest (ROIs) and their fluorescence time courses were extracted using constrained non-negative matrix factorization (CNMF). Components were retained based on spatial morphology, signal -to-noise ratio, and temporal activity and were visually inspected for segmentation quality. For each accepted ROI, fluorescence was background/ neuropil corrected, a time-varying baseline (F_0_) was estimated, and fractional fluorescence was calculated as ΔF/F_0_ = (F-F_0_)/F_0_). Traces were sampled at 1.18 Hz and used for subsequent measurements of transient amplitude, prominence, event frequency, inter-event interval, and network activity.

Separately, TIFF stacks were analyzed using a custom Python pipeline designed to detect both compact neuronal cell bodies and elongated projections. Pixelwise baseline fluorescence and temporal noise were estimated to identify spatially contiguous signals with sufficient intensity, signal-to-noise ratio, size, persistence, and positional stability. Cell contours were reconstructed by locally expanding stable soma-scale intensity cores, whereas projections were detected independently using fine-scale contrast and multiscale ridge analysis. Mean ROI fluorescence was corrected using the surrounding neuropil signal, normalized to a running 20th-percentile baseline, detrended, and lightly smoothed. Calcium events were accepted only when a connected multipixel signal persisted for at least three frames and remained spatially coherent and stationary. The resulting traces were used to calculate transient amplitude, prominence, signal-to-noise ratio, event frequency, event duration, and inter-event intervals.

Regions of interest corresponding to active cells/sites were identified within each recording. Individual regions of interest were considered measurements nested within an organoid and were not treated as independent biological replicates. The fluorescence signals were normalized to ΔF/F following baseline correction.

#### Quantitative PCR

RT-qPCR validation was performed using organoids generated from an independent differentiation. For each genotype, three replicates were analyzed, with each replicate comprising 6-6 pooled organoids and both independent isogenic clones represented. Total RNA was isolated from pooled organoids homogenized in Trizol™ reagent (Thermo Scientific) using the Direct-zol kit RNA extraction kit (Zymo Research). RNA was reverse transcribed into cDNA using the iScript Reverse Transcription Supermix (Bio-Rad). qPCR was performed using iTaq™ Universal SYBR Green Supermix (Bio-Rad) with previously published or commercially available primer sets^82, 88^ (see Key Resource Table).

## QUANTIFICATION AND STATISTICAL ANALYSIS

### Regulon Activity Statistical Evaluation

#### Comparative Metrics

Regulon activity patterns were compared across cell types, genotypes, and developmental time points, and visualized using UMAP embeddings, lineage-resolved trajectories, and Scanpy dot plots.

#### Analysis of Variance (ANOVA)

Regulon AUC scores were analyzed using a three-way ANOVA incorporating genotype, developmental time point, and cell type. Partial eta-squared (*η*^2^) values were calculated to quantify cell-type-associated effect sizes. Complete statistical results are cataloged in Supplementary **Table S2**.

### Base parameters statistical analysis

Event frequency and inter-spike intervals were calculated from detected calcium transients. Pairwise Spearman correlation coefficients were calculated among active regions within each organoid to assess coordinated activity. To compare network coordination among genotypes, pairwise correlations were summarized at the individual-organoid level, generating one biological-replicate measurement per organoid. Data are presented as mean ±SEM for biological replicates. One-way ANOVA with Bonferroni correction for multiple comparisons (alpha threshold = 0.05).

### Pairwise calcium activity correlation and hierarchical clustering

Cleaned ΔF/F traces from ROIs in the same organoid were compared across matched imaging frames using pairwise Spearman rank correlation (ρ). As a temporal control, one trace in each pair was reversed in time (frame N to frame 1), its correlation with the unchanged trace was recalculated, and this value was subtracted from the aligned correlation (Δρ = ρ_aligned − ρ_time-reversed). Reversal preserves each trace’s event count and temporal spacing while disrupting its original alignment with the other ROI; it does not constitute a complete correction for firing rate. ROIs were ordered by unsupervised average-linkage hierarchical clustering of the aligned correlations using 1 − ρ as the distance, and the same order was applied to heatmap rows and columns. The genotype blocks were displayed in WT, G191C, and G31A order, placing the stronger within-organoid correlations toward the upper left and weaker correlations toward the lower right. Self-correlations were excluded.

### Network synchrony analysis

Pairwise Spearman rank correlations were calculated between the cleaned ΔF/F traces of all active ROIs within each organoid, excluding self-correlations, and averaged to obtain one mean pairwise Spearman ρ per organoid. Organoids were treated as independent biological replicates (n = 15 per genotype). Organoid-level distributions were displayed as empirical cumulative step functions together with fitted logistic cumulative-distribution curves. Pointwise 95% confidence bands were generated from 5,000 bootstrap re-samples of whole organoids within each genotype, with the logistic curve refitted after each resampling. Genotype differences were assessed by one-way ANOVA followed by Bonferroni-adjusted pairwise comparisons (α = 0.05); group medians and adjusted P values are reported in the panel.

### Calcium-wave propagation analysis

Temporally coordinated calcium events were identified across active somatic ROIs, with inactive or structural ROIs excluded. For each representative wave, the arrival time at every responsive ROI was defined from the detected calcium-event peak relative to the first responding ROI. ROI centroid coordinates were restricted to a common 512 × 512-pixel field, and relative arrival times were spatially interpolated within the convex hull of the responsive ROIs to generate isochrone maps. Color indicates arrival delay in seconds, while normalized white vectors represent the local direction of increasing arrival time and not velocity magnitude. Propagation speed was estimated from the spatial–temporal arrival-time gradient and reported in pixels per second. WT, G31A, and G191C are representative examples illustrating fast, slow, and intermediate propagation, respectively.

### Calcium-event phase analysis

To quantify the timing of cellular calcium events relative to population activity, a leave-one-ROI-out population signal was generated for each active ROI from the standardized ΔF/F traces of all other active ROIs in the same organoid. The population signal was band-pass filtered at 0.01–0.20 Hz using a zero-phase, second-order Butterworth filter, and its instantaneous phase was calculated from the angle of the Hilbert analytic signal. Each validated calcium-event peak, excluding the first and last 20 s of each recording, was assigned the corresponding population phase. Phase-bin fractions were calculated separately for each organoid and averaged with equal weighting across 15 organoids per genotype to generate the rose plots. Circular mean angle and mean resultant length were calculated per organoid to quantify preferred phase and phase concentration, respectively. Phase locking was evaluated against 100 ROI-wise circular-shift surrogates, and organoids, not individual ROIs or events, were treated as biological replicates.

## RESOURCE AVAILABILITY

### Lead contact

- Requests for further information and resources should be directed to and will be fulfilled by the lead contact, Derrick J. Morton.

### Materials availability

- All unique/stable reagents generated in this study are available from the lead contact upon execution of a completed materials transfer agreement.

### Data and code availability

- scRNA-seq data have been deposited at NCBI GEO at accession number [GSE329516] and are publicly available as of the date of publication.
- All scripts used to analyze this dataset are available on https://github.com/DMortonlab/Cerebellar-Organoid-scRNA-seq.
- Any additional information required to reanalyze the data reported in this paper is available from the lead contact upon request.

## ACKNOWLEDGMENTS

This work was supported by a National Institutes of Health R01 grant (NS131620) and an Alfred P. Sloan Research Fellowship in Neuroscience award (FG-2023-20698) to D.J.M.; a National Institutes of Health DP1 grant (NIMH132709) to K.D. N.A.B. was supported by the Ruth L. Kirschstein National Research Service Award (F31NS139645) and a University of Southern California WiSE Scholarship. We thank Dr. Giorgia Quadrato and her laboratory at the University of Southern California for their guidance on cerebellar organoid generation.

## AUTHOR CONTRIBUTION

Conceptualization, N.A.B. and D.J.M.; methodology, N.A.B, H.K., and D.J.M, Investigation, N.A.B., H.K., R.E.K., J.J.G., M.J.W., E.N.T., V.L., B.B., writing—original draft, N.A.B., and D.J.M.; writing—review & editing, N.A.B., A.E.S, K.D., B.B., H.K., D.J.M.,; funding acquisition N.A.B., K.D., D.J.M.; resources, D.J.M; supervision, D.J.M.

## Supplemental Figure Legends

**SFig. 1:**
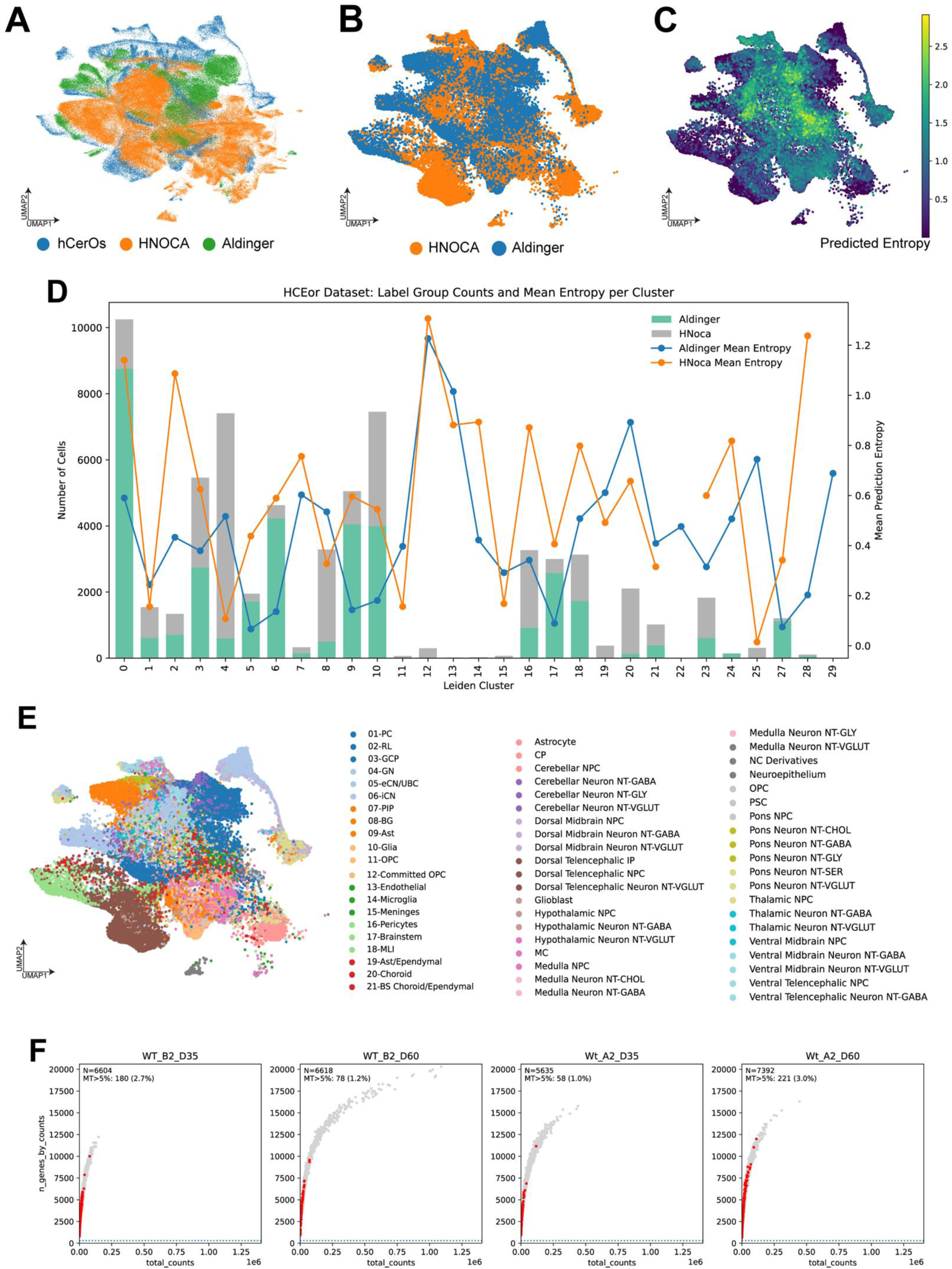
Reference atlas mapping and quality control of hCerO scRNA-seq datasets. **(A)** Integrated UMAP of hCerO, HNOCA, and Aldinger reference datasets. **(B)** UMAP of HNOCA and Aldinger reference cells (C) UMAP showing predicted cell type annotation entropy **(D)** Reference cell counts and mean prediction entropy by Leiden cluster. **(E)** UMAP showing hCerO and reference cell type annotation **(F)** Quality control plots showing detected genes, total transcript counts, and mitochondrial transcript percentages in independent EXOSC3WT biological replicates at D35 and D60.

**SFig. 2:**
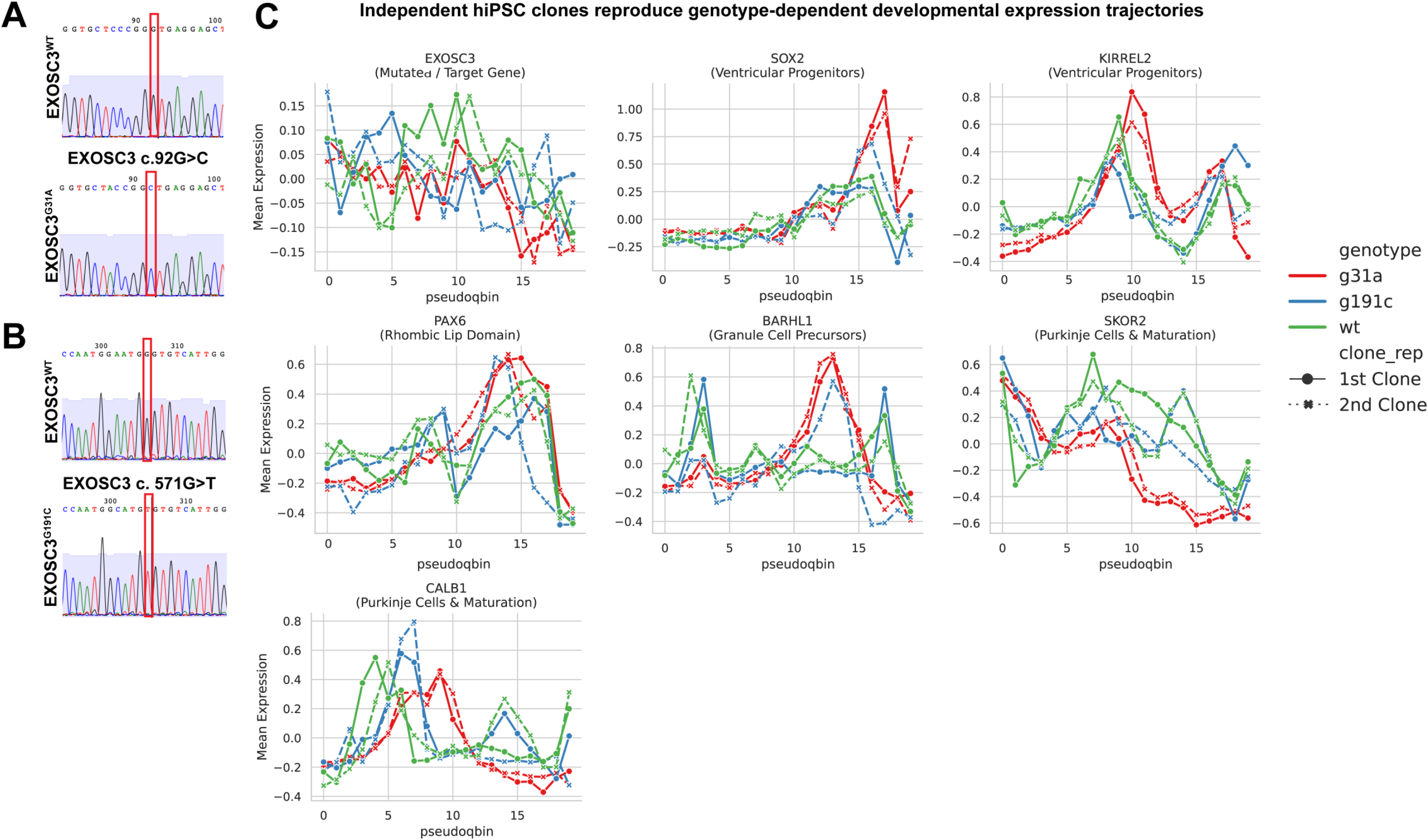
Validation of EXOSC3 mutations and reproducibility of developmental expression trajectories. **(A-B)** Sanger sequencing confirms the engineered EXOSC3 c. 92G>C (G31A) and c. 571G>T (G191C) mutations in isogenic hiPSC lines. Red boxes indicate the edited nucleotides. **(C)** Replicate-resolved expression trajectories of EXOSC3, SOX2, KIRREL2, PAX6, BARHL1, SKOR2, and CALB1 along Purkinje-lineage pseudotime in WT, G31A, and G191C hCerOs. Solid and dashed lines represent two independently derived hiPSC clones per genotype.

**SFig. 3.**
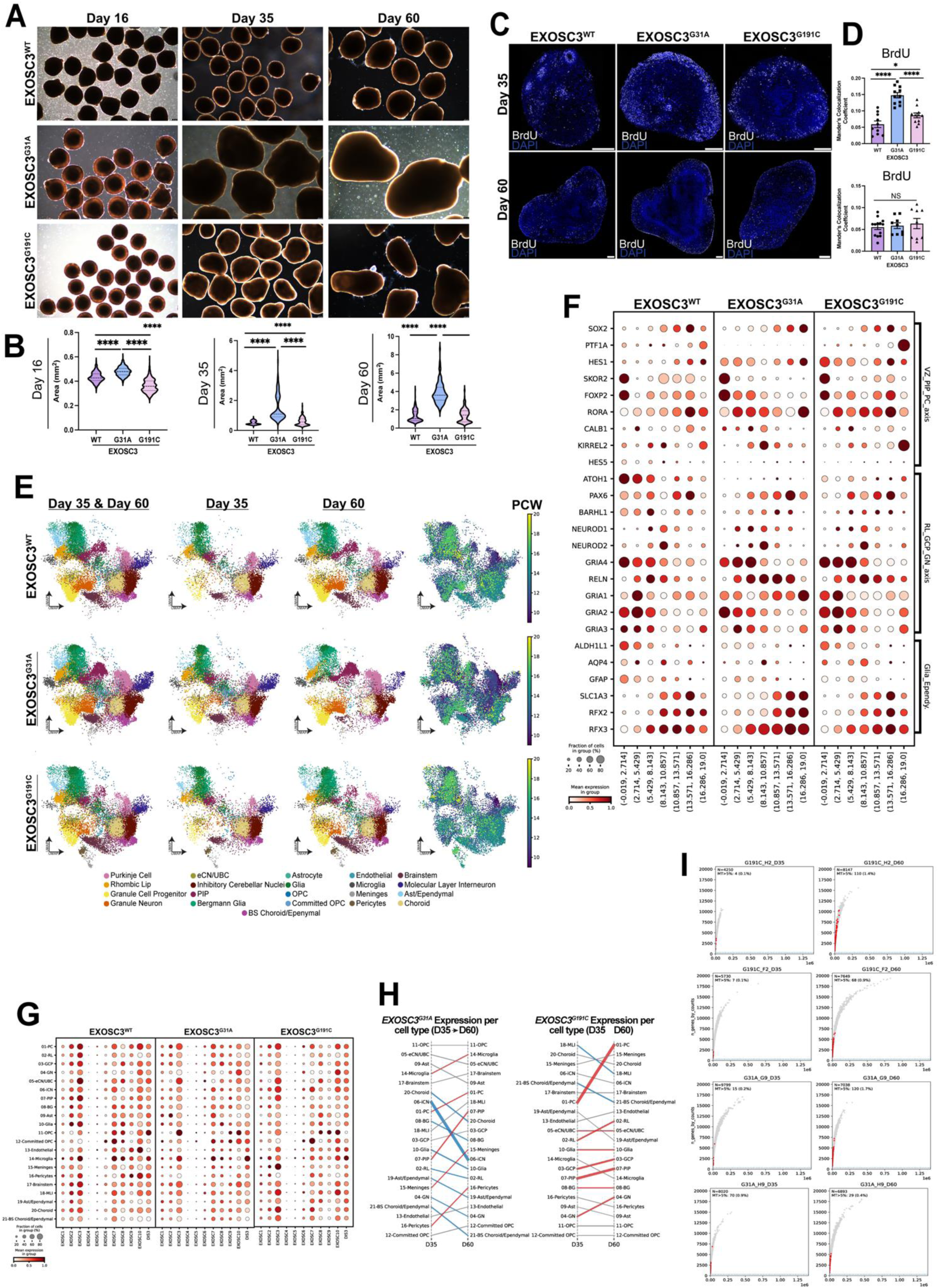
Characterization of EXOSC3 mutant cerebellar organoids and scRNA-seq datasets. **(A)** Representative brightfield images of EXOSC3^WT^, EXOSC3^G31A^, and EXOSC3^G191C^ hCerOs at D16, D35, and D60. **(B)** Quantification of organoid areas at each developmental time point. **(C-D)** Representative BrdU immunostaining and quantification of proliferating cells at D35 and D60. **(E)** UMAPs showing cell type distributions at D35 and D60 and predicted developmental age in postconception weeks (PCW) for each genotype. **(F)** Dot plots showing expression of canonical cerebellar lineage markers in WT and mutant hCerOs. **(G)** Dot plots showing RNA exosome subunit expression among cell populations by genotype. **(H)** Changes in EXOSC3 expression among cell populations between D35 and D60 in G31A and G191C organoids. **(I)** Single-cell RNA-seq quality control showing detected genes, total transcript counts, and mitochondrial transcript percentages in independent mutant biological replicates. Statistical significance: ****p<0.0001; NS, not significant.

**SFig 4.**
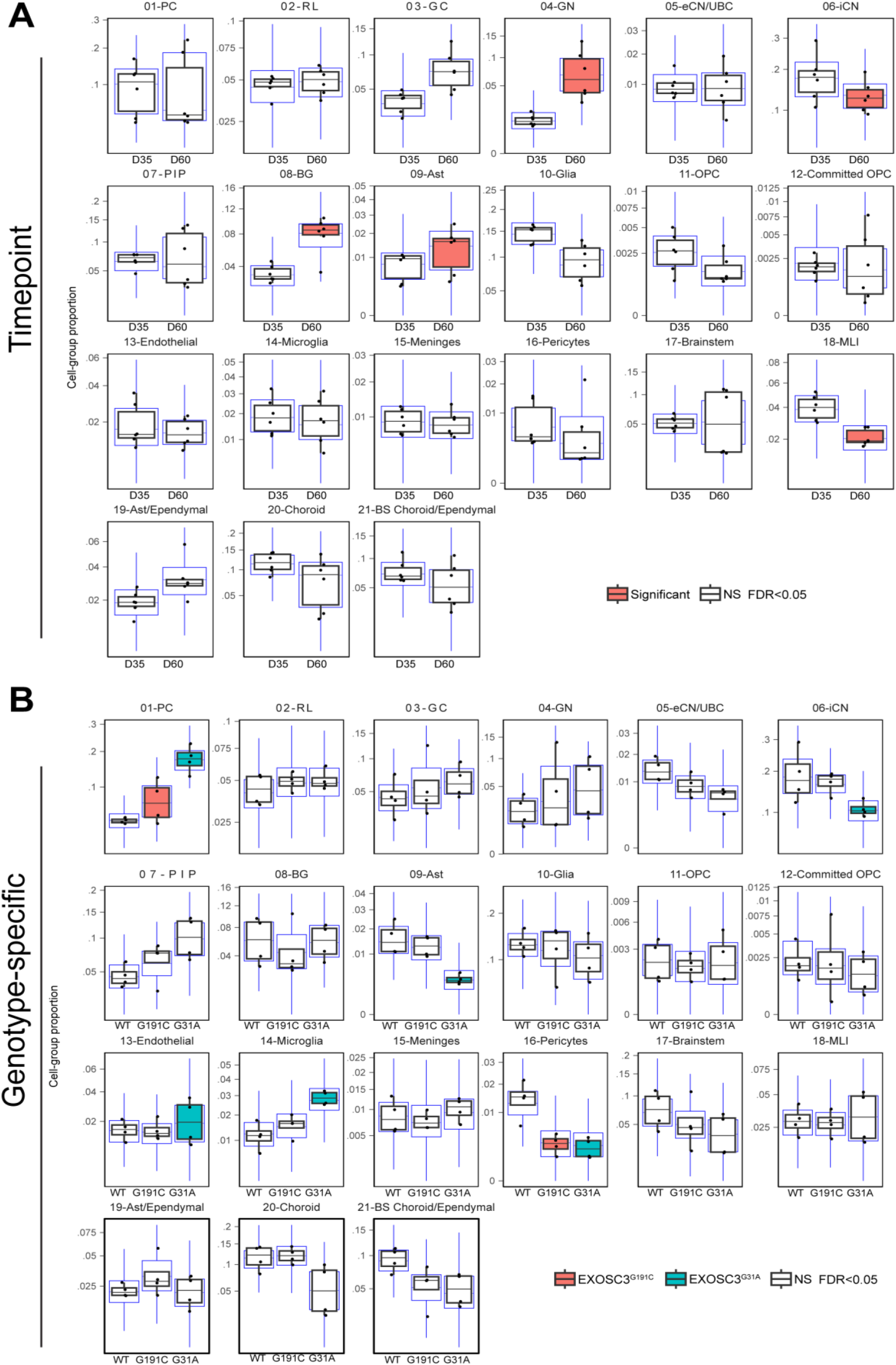
Developmental and genotype-dependent changes in cerebellar cell type composition. **(A)** Changes in cell type proportions between D35 and D60 in hCerOs. **(B)** Genotype-dependent differences in cell type proportions among EXOSC3^WT^, EXOSC3^G31A^, and EXOSC3^G191C^ hCerOs. Differential abundance was assessed using sccomp model. Box plots show cell type proportions, with individual observations indicated by points. Colored boxes indicate significant differences (FDR<0.05); white boxes indicate nonsignificant differences.

**SFig. 5:**
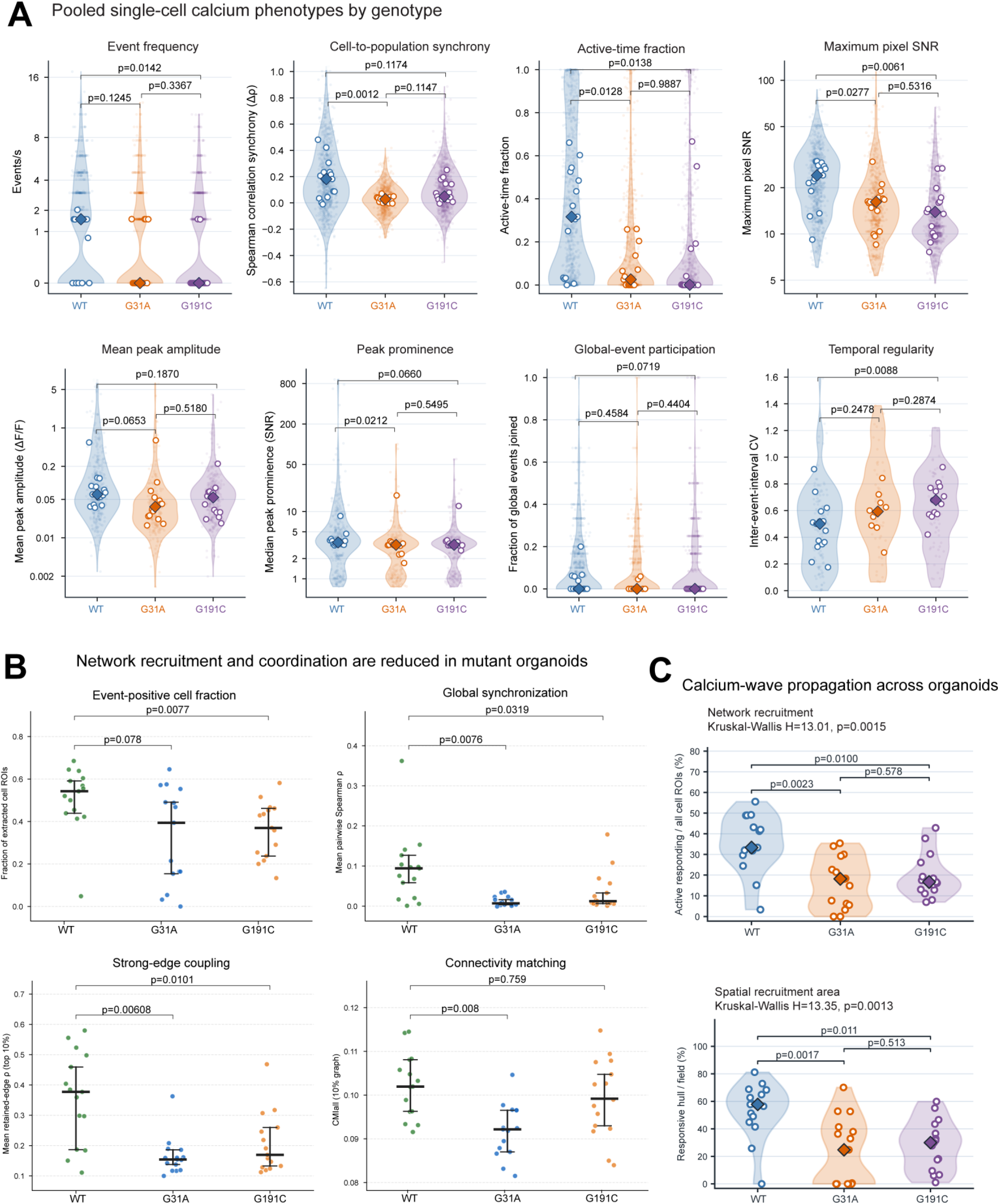
EXOSC3 variants alter spontaneous calcium activity and neuronal network coordination. **(A)** Violin plots showing pooled single-cell calcium phenotypes in EXOSC3^WT^, EXOSC3^G31A^, and EXOSC3^G191C^ hCerOs, including event frequency, cell-to-population synchrony, active time fractions, maximum pixel signal-to-noise ratio (SNR), peak amplitude, peak prominence, global event participation, and temporal regularity. Open circles represent organoid-level values. p-values are indicated. **(B)** Organoid-level quantification of event-positive cell fraction, global synchronization, strong-edge coupling, and connectivity matching (n =15 organoids/genotype. Points represent individual organoids; horizontal bars indicate group summaries and error intervals. p-values are indicated. **(C)** Quantification of calcium-wave network recruitment and spatial recruitment area among genotypes. Violin plots show organoid-level distributions. Kruskal-Wallis and pairwise adjusted p-values are indicated.

**SFig. 6:**
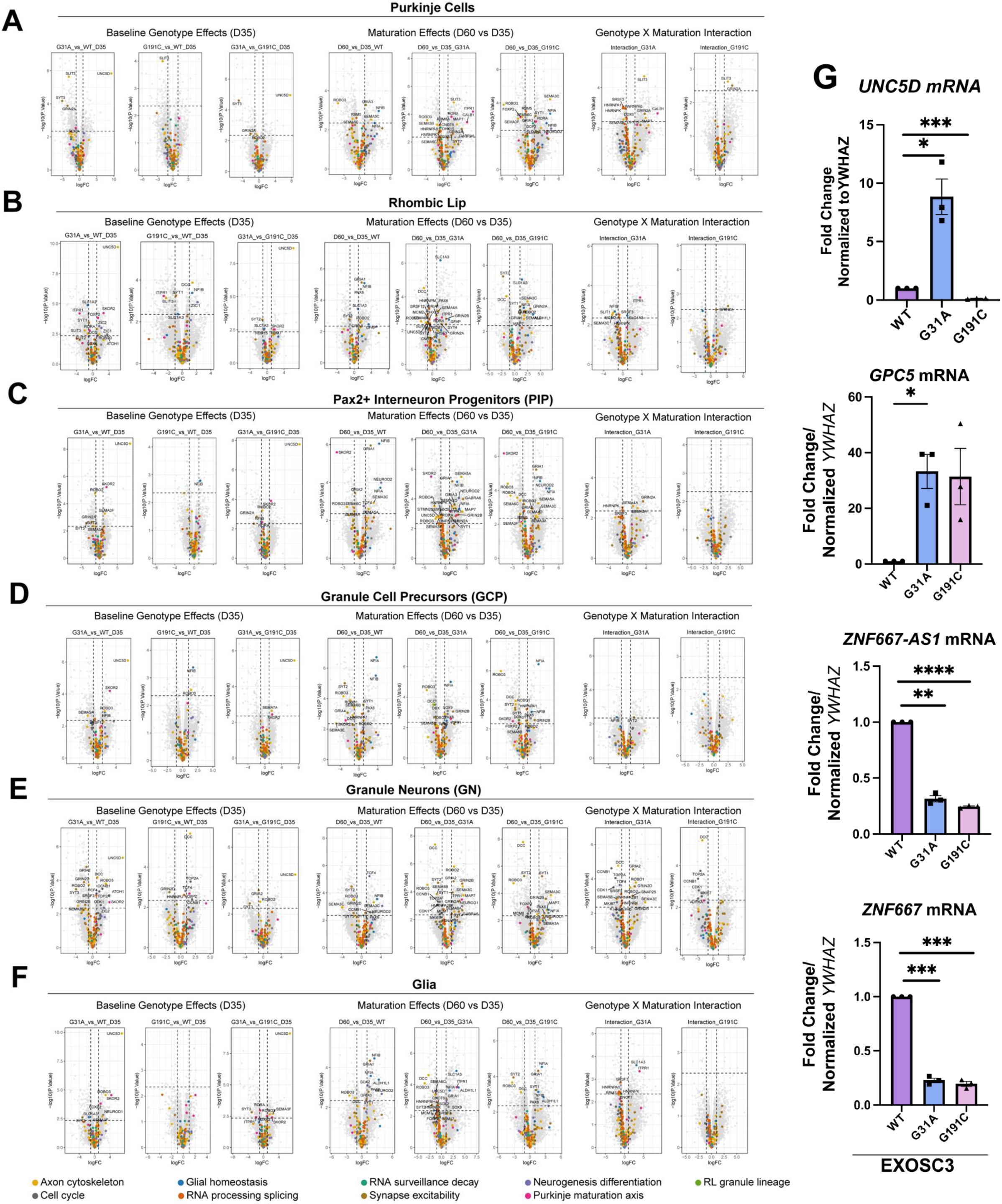
Lineage-specific transcriptomic changes associated with EXOSC3 dysfunction during cerebellar development. **(A-F)** Volcano plots showing differential gene expression in Purkinje cells (**A**), rhombic lip (**B**), PAX2+ interneuron progenitors (**C**), granule cell precursors (**D**), granule neurons (**E**), and glia (**F**). Comparisons include baseline genotype effects at D35 (G31A vs. WT, G191C vs. WT, and G31A vs. G191C), maturation-associated changes (D60 vs. D35 within each genotype), and genotype-by-maturation interactions. Colored points indicate genes associated with neuronal differentiation, axon organization, synaptic function, RNA processing, cell cycle regulation, and cerebellar lineage development, as indicated in the legend. **(G)** RT-qPCR validation of *UNC5D, GPC5*, *ZNF667-AS1*, and *ZNF667* expression in D35 WT, G31A, G191C hCerOs. Expression was normalized to YWHAZ and presented relative to WT. Statistical significance was assessed using unpaired two-tailed Welch’s t-tests.

**SFig. 7:**
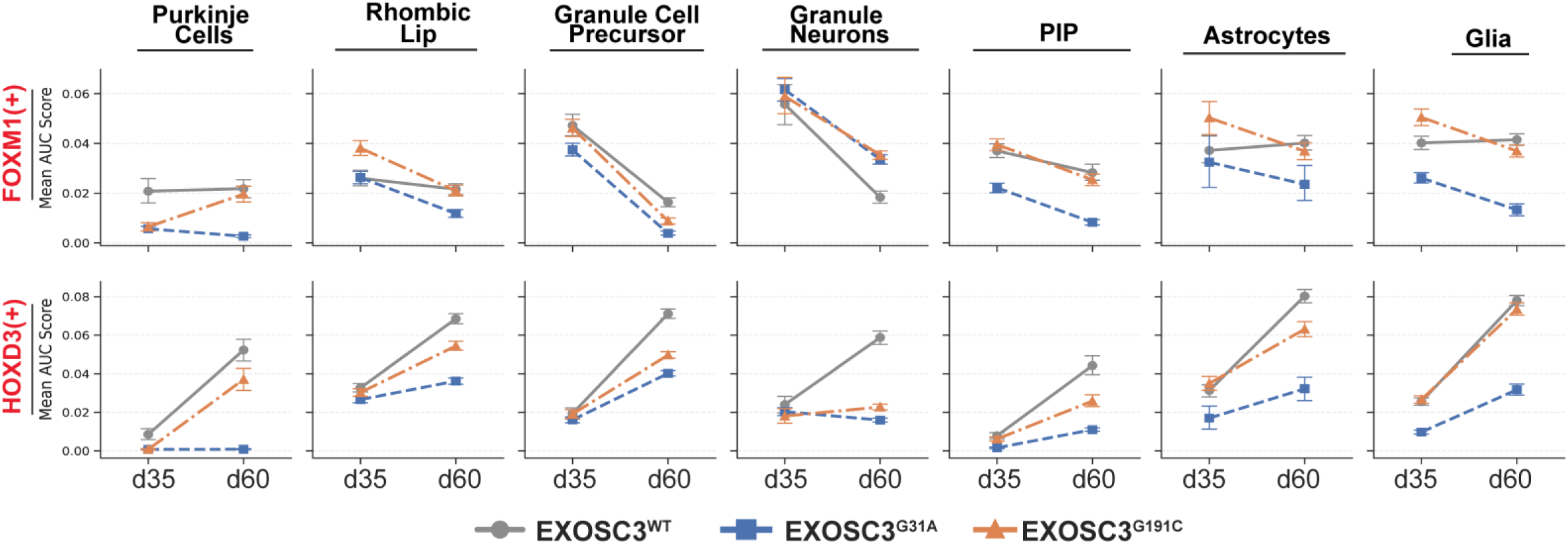
Developmental changes in FOXM1- and HOXD3-associated regulon activity in *EXOSC3* mutant cerebellar organoids. Mean SCENIC regulon activity (AUCell AUC scores) for FOXM1(+) and HOXD3(+) and D35 and D60 in Purkinje cells, rhombic lip, granule cell precursors, granule neurons, PAX2+ interneuron progenitors, astrocytes, and glia. Error bars represent SEM. Global statistical analyses identified significant genotype-, developmental timepoint-, and cell type-associated interaction effects (**Table S2**).

